# Neutrophil-driven cardiac damage during invasive *Streptococcus pneumoniae* infection is regulated by CD73

**DOI:** 10.1101/2022.12.07.519499

**Authors:** Manmeet Bhalla, Vijay R. Ravi, Alexsandra Lenhard, Essi Y. I. Tchalla, Jennifer K Lang, Elsa N. Bou Ghanem

## Abstract

*Streptococcus pneumoniae* (pneumococcus)-induced cardiac events are one of the life-threatening infection outcomes of invasive pneumococcal disease. *S. pneumoniae* has the ability to invade the myocardium and damage cardiomyocytes, however the contribution of the immune response during this process is not fully understood. We previously found that polymorphonuclear cells (PMNs) are crucial for host defense against *S. pneumoniae* lung infection and that extracellular adenosine (EAD) production, by exonucleosidases CD39 and CD73, controlled the anti-bacterial functions of these cells. The objective of this study was to explore the role of PMNs and the EAD-pathway in host cardiac damage during invasive pneumococcal infection. Upon intra-peritoneal (*i*.*p*.) injection with invasive *S. pneumoniae* TIGR4 strain, hearts of C57BL/6 mice showed an increased influx of PMNs as determined by flow cytometry. However, the increased PMN numbers failed to contain the bacterial burden in the heart and showed positive correlation with serum levels of the cardiac damage marker Troponin-1. Influx of PMNs into the heart was associated with constant presence of neutrophil degranulation products in the cardiac tissue. Depletion of PMNs prior infection reduced pneumococcal burden in the heart and lowered the Troponin-1 levels thus, indicating their role in cardiac damage. While exploring the mechanisms underlying the damaging PMN response, we found that by 24hpi, there was a significant reduction in the expression of CD39 and CD73 on cardiac PMNs. The role of CD73 in regulating cardiac damage was tested *in vivo* using CD73^-/-^ mice which had significantly higher bacterial burden and cardiac damage compared to wild type mice despite similar PMN numbers. The role of CD73 expression on PMNs was also tested *ex vivo* using the HL-1 cardiomyocyte cell line which upon *S. pneumoniae* infection, showed increased cell death in presence of CD73^-/-^ PMNs. Our findings have identified a detrimental role for PMNs in cardiac damage during invasive pneumococcal infection that is in part driven by reduced expression of EAD-producing enzymes in late disease stages.

## 1. Introduction

*Streptococcus pneumoniae* (pneumococcus) are Gram-positive opportunistic pathogens that typically reside asymptomatically in the upper respiratory tract of a healthy individual. Under certain conditions, these bacteria can cause serious and life-threatening infections [1, 2], resulting in over a million deaths annually on the global scale [1]. The infections caused by *S. pneumoniae* predominantly include pneumonia but certain stains of pneumococci are capable of causing invasive pneumococcal disease (IPD) leading to bacteremia, meningitis and endocarditis [1]. In recent years, pneumococcal-induced cardiac events have emerged as one of the most serious and life-threatening infection outcomes [3, 4] leading to major adverse cardiac events (MACE). These include myocardial infarction, new or worsening of arrhythmias, collagen deposition in individuals treated with antibiotics and congestive heart failure with high mortality rate [3-5]. In fact, the risk of a major cardiac event can persist up to 10 years in individuals treated for pneumonia [6]. Central to pneumococcal-induced adverse cardiac events is the ability of the bacterium to translocate from the circulation into the myocardium, induction of cardiomyocyte death through the release of bacterial factors and formation of bacteria-filled cardiac microlesions culminating into impaired electrophysiology and contractile ability of the heart muscle cells [7-9]. While bacterial factors driving cardiac damage are starting to be elucidated [8], the role of the immune response in cardiac damage during invasive pneumococcal infection has not been characterized.

We and others previously showed that Polymorphonuclear leukocytes (PMNs), also known as neutrophils, can shape the course of disease following pulmonary *S. pneumoniae* infection [10-13]. The presence of PMNs is important for adequate control of bacterial burden during the initial phase of lung infection [10, 12]. Depletion of PMNs prior to intratracheal challenge with *S. pneumoniae* leads to higher bacterial numbers in the lungs and increased host mortality [10] thus, demonstrating the importance of these cells in host resistance to pulmonary infection early on. However, we previously found that the phenotype and anti-bacterial response of PMNs is altered during the course of pneumococcal pneumonia. In fact, depletion of PMNs 18 hours following pulmonary challenge with *S. pneumoniae* reduced bacterial numbers in the lungs. Therefore, in later phases of pneumococcal pneumonia, PMNs become detrimental to host survival [10]. Most of the data regarding the host immune response against pneumococcal infection is focused in the lung environment [14, 15], given that pneumonia is the most common disease outcome. However, there is a limited information about how the host responds during pneumococcal invasion of the cardiac tissue. A prior study showed that *S. pneumoniae*, once in the heart, release pneumolysin which kills infiltrated macrophages at the site of bacterial microlesions [8] leading to the blunting of the host immune response to the infection. Although reported to be detected in the heart during pneumococcal endocarditis [8], the role of PMNs in host susceptibility to cardiac infection and cardiomyocyte damage during IPD has not been investigated.

One of the host pathways that regulate PMN responses during pneumococcal pneumonia is the extracellular adenosine (EAD)-pathway [16]. EAD is produced by sequential conversion of ATP, the level of which increases during tissue injury, by two exonucleosidases CD39 and CD73 [16, 17]. EAD can then act and signal through any of the four downstream G-coupled protein receptors, A1, A2A, A2B and A3 [17]. We previously showed the importance of CD73 in PMN pulmonary influx and resolution, and the role of CD73 in PMN-mediated bacterial killing during pneumococcal pulmonary infection [10, 13]. EAD plays a critical role in regulating cardiac functions and heart electrophysiology [18]. Adenosine receptors downstream of CD73 are important regulators of heart rate, myocardial contractility and coronary flow [18]. However, the role of CD73 in cardiac damage driven by infection has not been explored.

In this study, we explored the role of PMNs and CD73 in cardiac damage during invasive pneumococcal infection. We found that PMNs migrated into the heart following pneumococcal challenge and that their influx into the cardiac tissue positively correlated with cardiomyocyte damage. Depletion of PMNs prior challenge ameliorated cardiomyocyte injury and reduced the bacterial burden in the cardiac tissue. We found that during the late stage of IPD, increased bacterial burden and exacerbated cardiomyocyte damage coincided with reduced CD73 expression on PMNs. We tested the role of CD73 *in vivo* and found that CD73^-/-^ mice showed significantly higher bacterial burden and increased cardiomyocyte damage compared to WT controls. CD73 expression on PMNs was important to prevent cardiac damage as *in vitro* HL-1 cardiomyocyte showed decreased cell viability in presence of CD73^-/-^ PMNs compared to the wild-type controls upon infection. These findings reveal a novel role for PMNs in host susceptibility to cardiac damage during invasive *S. pneumoniae* infection and identifies CD73 expression on these cells to be important for host tissue protection.

## 2. Results

### 2.1 PMNs influx into the cardiac tissue following *S. pneumoniae* systemic challenge

To understand the role of PMNs in cardiac tissue damage during invasive pneumococcal infection, we first studied the kinetics of PMN influx in the heart during the course of infection with *S. pneumoniae*. To do so, we used a murine systemic model of infection established by the Orihuela group and shown to results in pneumococcal infection and damage of the myocardium [7, 8, 19, 20]. C57BL/6 mice were infected intra-peritoneally (I.P.) with 10^3 colony-forming units (CFU) of *S. pneumoniae* TIGR4 strain and PMN influx and bacterial burden were monitored at 6-, 12- and 24 hours post-infection (hpi) in the blood and heart. Bacterial challenge through intra-peritoneal route was done to synchronize the infection, as previously shown [8, 20]. Infection with *S. pneumoniae* resulted in an increase in PMN (Ly6G^+^) numbers in the blood, as determined by flow cytometry, as early as 6hpi and peaked at 24hpi (Fig 1B), the time point when mice got severely sick and had to be euthanized. Quantification of bacterial numbers in the blood showed a linear increase in bacterial burden (Fig 1D) as the infection progressed despite increase in the PMN numbers (Fig 1B).

**Fig 1.**
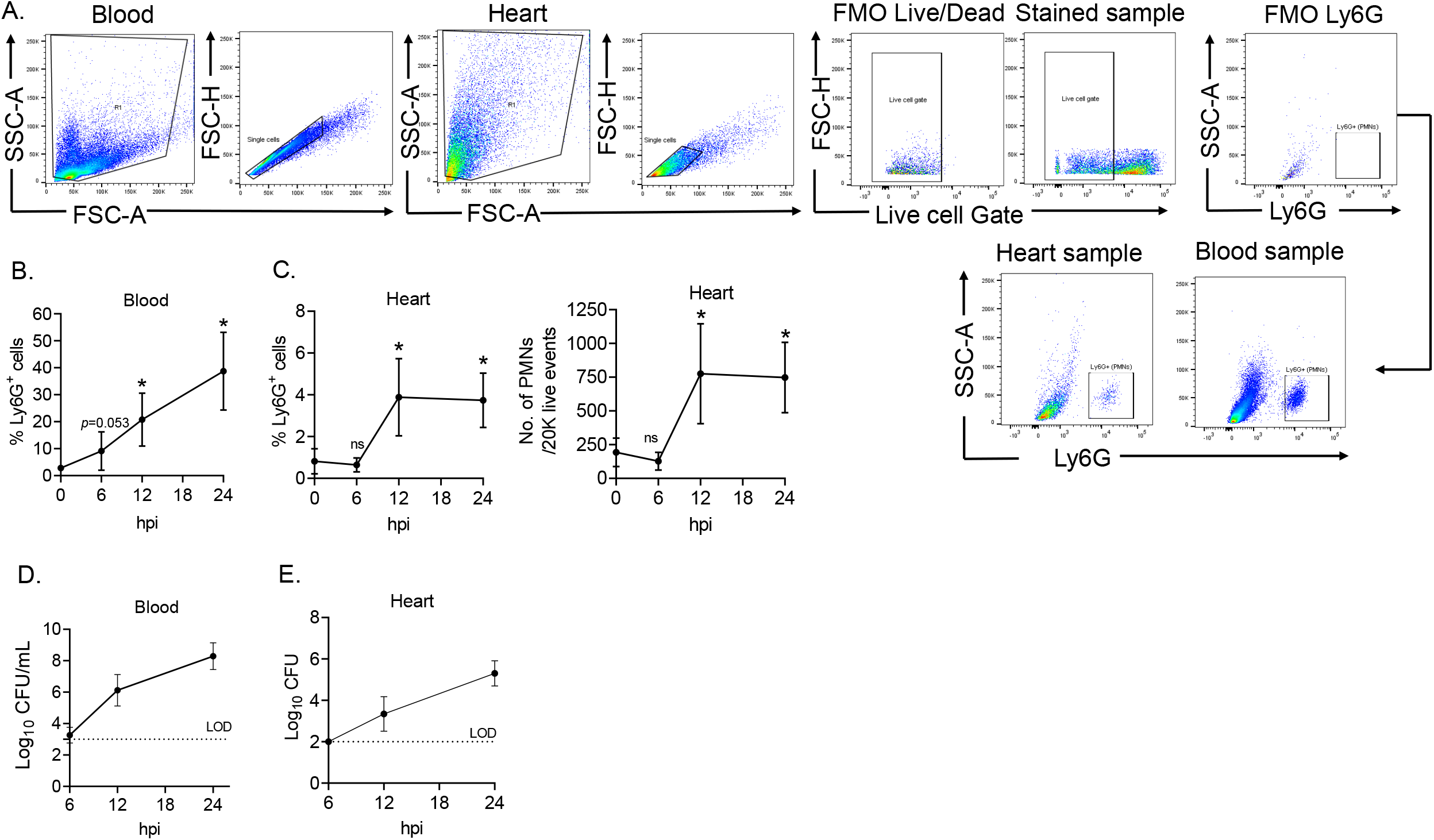
Infection with *S. pneumoniae* TIGR4 strain triggers influx of PMNs into the host cardiac tissue. WT (C57BL/6) mice were injected infected (I.P) with 10^3 CFU of *S. pneumoniae* TIGR4 or mock infected with PBS. The mice were euthanized at 6-, 12- and 24hpi to harvest blood and heart to enumerate bacterial burden and PMN numbers. (A) Gating strategy showing PMN population (Ly6G^+^) in the blood and heart. (B-E) The % PMNs in blood (B) %PMNs and PMN numbers per 20K live cells in heart (C) along with the respective bacterial burden (D-E) are shown. Data shown (B-D) are from a total of 6 mice per group/time point. CFU data (D-E) are pooled from n=6 mice for 6- and 24hr time points and 12 mice for 12hr time point. Asterisks indicate significant differences as calculated by unpaired Student’s t-test comparing the infected group to the baseline, where *p*<0.05 is considered as significant.

Flow cytometric analysis of single-cell suspension of the whole cardiac tissue showed no increase in PMN numbers at 6hpi (Fig 1C) despite increase in blood PMN numbers. However, the cardiac influx of PMNs increased and peaked at 12hpi (Fig 1C). Use of cell-viability dye ensured that the observed increase in PMN numbers was not an artifact of analyzing dead cells (Fig S1). Interestingly, the number of PMNs in the cardiac tissue did not show a further increase between 12- and 24hpi (Fig 1C) despite an approximately 2-fold increase in their numbers in the blood during the same time period (Fig 1B) and an increased bacterial presence in the heart (Fig 1E). Enumeration of bacterial burden in the heart showed a significant positive correlation with the pneumococcal numbers in the blood (Fig S2), as shown previously [20].

To test if the cardiac influx of PMNs is bacterial strain specific, mice were challenged intraperitoneally with 10^3 CFU of D39, another invasive strain of *S. pneumoniae* [20]. As observed with TIGR4, we saw the presence of both D39 and PMNs (Fig S3) in the cardiac tissue of these mice over the course of infection indicating that the invasive strains of *S. pneumoniae* that translocate into the myocardium induce the recruitment of PMNs into the heart.

To confirm if pulmonary infection with *S. pneumoniae*, the route which mimics the primary manifestation of pneumococcal infection, recapitulates the above findings, mice were intra-tracheally challenged with 2.5 × 10^5 CFU of TIGR4 strain. At 24hpi, we detected the presence of bacteria in the lungs, circulation and heart (Fig 2C). PMN numbers also increased in the heart of these mice (Fig 2B) along with increased serum Troponin-I levels (indicative of cardiomyocyte injury) (Fig 2D). Taken together, these findings indicate that invasive infection with *S. pneumoniae* results in PMNs influx into the cardiac tissue.

**Fig 2.**
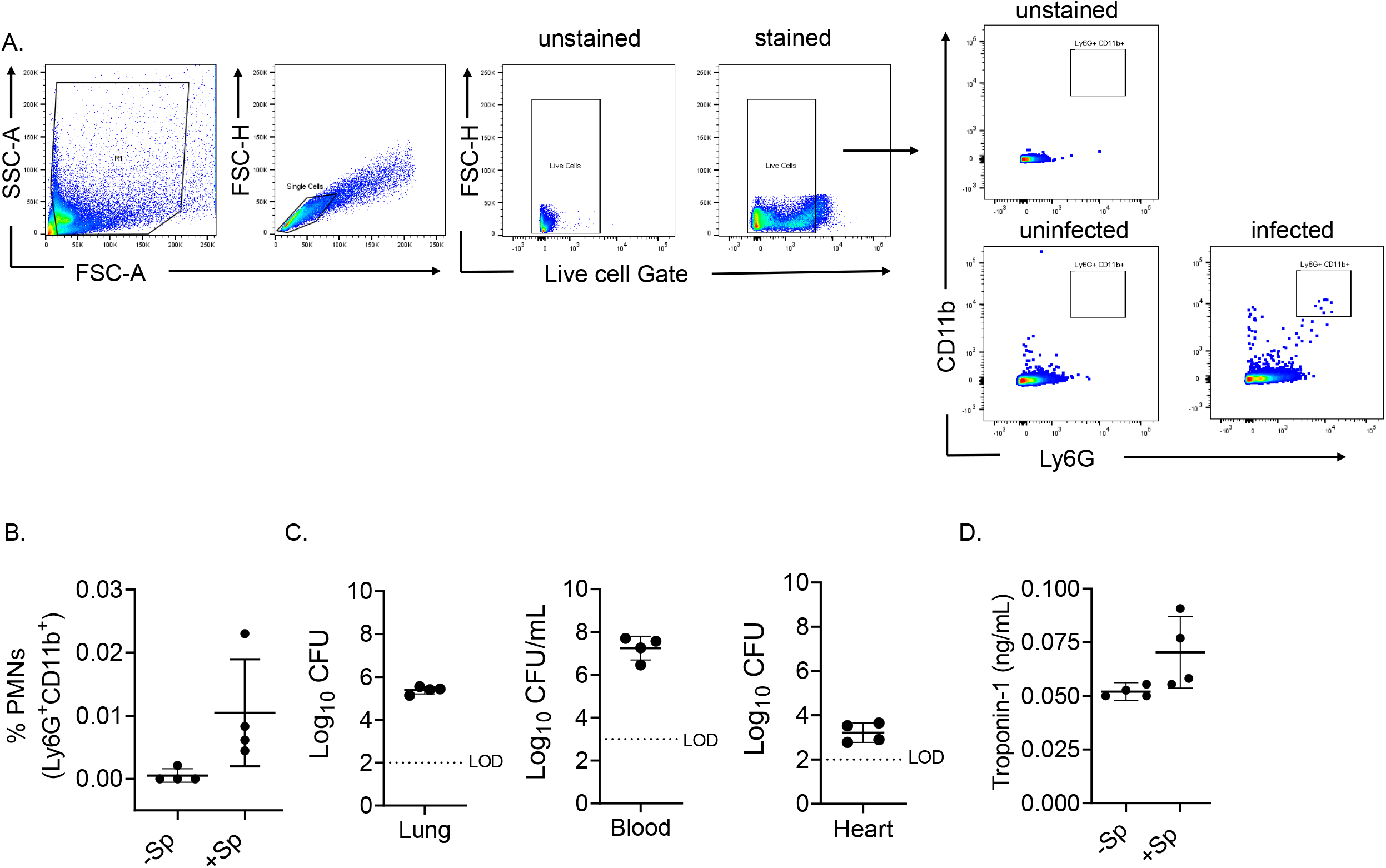
Intra-tracheal route of infection with *S. pneumoniae* results in infection of the host cardiac tissue. WT (C57BL/6) mice were infected intra-tracheally (I.T) with 2 × 10^5 CFU of *S. pneumoniae* TIGR4 and heart tissue was harvested 24hpi to detect PMN influx and bacterial presence. (A) Gating strategy showing PMN population (Ly6G^+^CD11b^+^) in the heart. (B) Enumeration of PMNs in the heart. (C) Bacterial load was enumerated in lungs, blood and heart. (D) Troponin levels in the sera samples. Data shown are from n = 4 mice per group in both uninfected and infected groups where every circle indicates one mouse. Data for serum Troponin-I was analyzed through Student t-test with Welch correction. LOD: Limit of detection

### 2.2 Influx of PMNs in the cardiac tissue correlates with myocardial damage

We next wanted to determine the effect of PMN influx on the host myocardial tissue. Serum samples were collected at 6-, 12- and 24hpi from mice infected intraperitoneally with 10^3 CFU of *S. pneumoniae* TIGR4 strain and tested for troponin levels as a marker of cardiomyocyte injury [7-9]. Mice challenged with PBS served as uninfected controls. We saw a non-significant increase in troponin-I levels at 12hpi (Fig 3A) and a significant increase in troponin-I levels at 24hpi (Fig 3A), indicating an increased cardiomyocyte damage with the progression of the disease. Interestingly, a positive correlation (r=0.52 and p=0.05) was observed between PMN presence in the heart and the cardiac tissue damage (Fig 3B) in infected mice, hinting towards a potential role of PMN in tissue injury.

**Fig 3.**
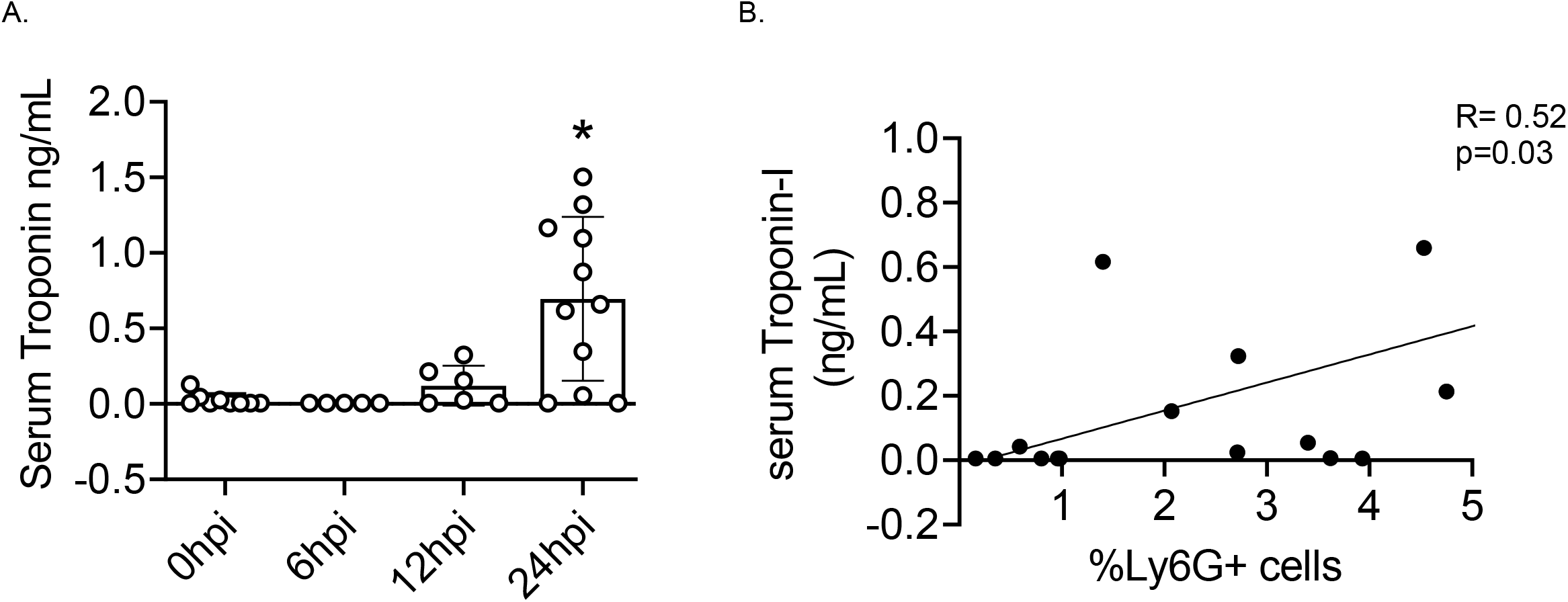
Influx of PMNs into the cardiac tissue during invasive pneumococcal infection is associated with increased cardiomyocyte damage. WT (C57BL/6) mice were infected intra-peritoneally (I.P) with 10^3 CFU of *S. pneumoniae* TIGR4 and mice were euthanized at 6-, 12- and 24hpi to harvest blood for Troponin-I measurement or heart tissue to enumerate PMN numbers in the heart. Data shows the (A) levels of serum Troponin-I levels at the indicated time points and (B) correlation between the troponin-I levels and %PMNs in the infected host as calculated by Pearson correlation test. Data shown are pooled from 6-11 mice per group where every circle indicates one mouse. Asterisks indicate significant differences as calculated by One-way ANOVA followed by Tukey’s test where *p*<0.05 is considered as significant.

### 2.3 Depletion of PMNs results in reduced bacterial numbers and less myocardial damage following systemic *S. pneumoniae* infection

To experimentally address the role of PMNs during invasive pneumococcal infection, mice were depleted of PMNs with intra-peritoneal (i.p.) injections of anti-Ly6G antibody (clone IA8) [10] a day prior and on the day of i.p. infection with 10^3 CFU of *S. pneumoniae* TIGR4 strain. Treatment with the anti-Ly6G antibodies resulted in >95% depletion of circulating and cardiac PMNs (Fig S4B) compared to isotype-treated controls. Depletion of PMNs did not affect the overall numbers of macrophages (Fig S4C) or dendritic cells (Fig S4D) in the cardiac tissue. The effect of PMN depletion on bacteremia, bacterial burden in the heart and serum troponin levels was determined and compared between the groups at 24hpi. When comparing the bacterial numbers in the blood, interestingly we did not observe any significant difference in the bacterial load between the two groups (Fig 4A), suggesting that during the late stage of invasive pneumococcal infection, PMN presence is not contributing to host resistance against *S. pneumoniae*.

**Fig 4.**
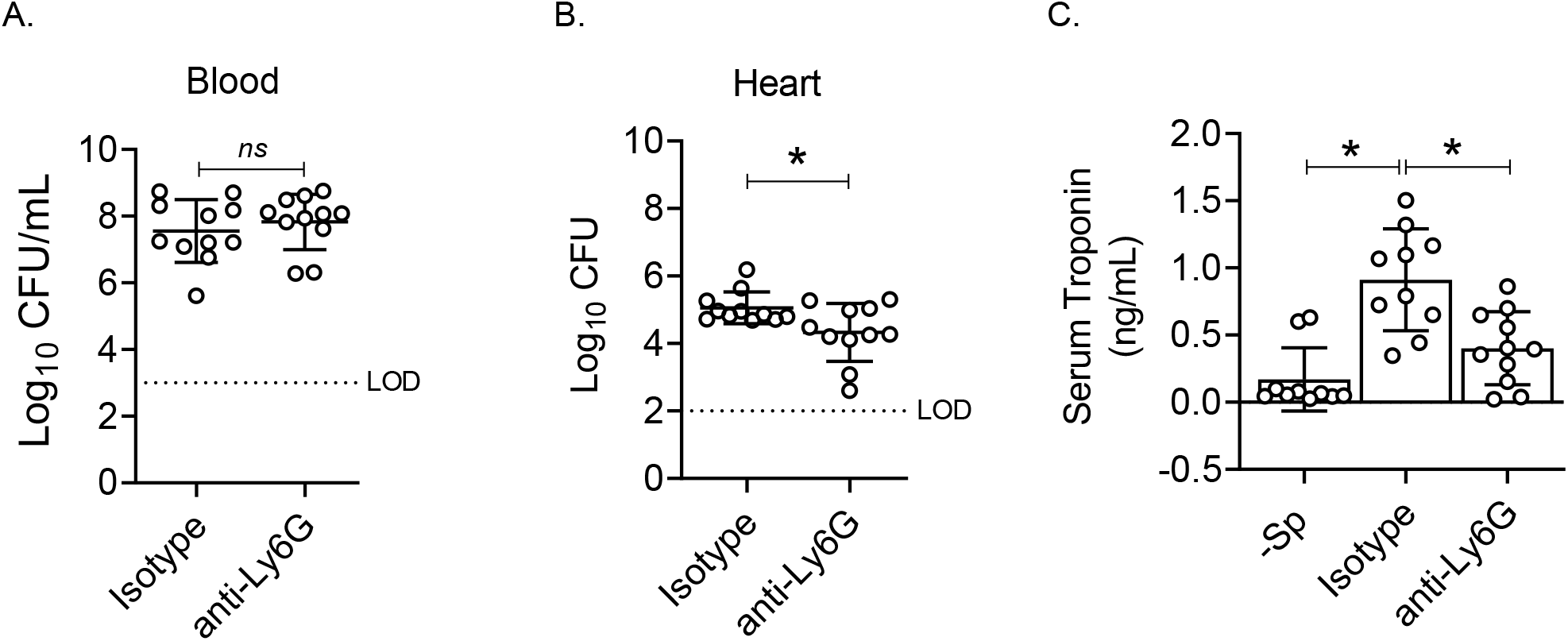
PMN depletion during systemic pneumococcal infection protects against cardiac damage. WT (C57BL/6) mice were injected intra-peritoneally (I.P) with either anti-Ly6G (clone 1A8) antibody or isotype control. At 18 hours later, the mice were injected with another dose of anti-Ly6G antibody and infected (I.P) with 10^3 CFU of *S. pneumoniae* TIGR4 or mock infected with PBS. The mice were then euthanized 24hpi and (A) bacterial burden in the blood and (B) the heart was enumerated and (C) the levels of Troponin-I in the sera samples were quantified. Data shown are pooled from a total of 9-11 mice per group where every circle indicates one mouse. Asterisks indicate significant differences as calculated by Student’s t-test with *p*<0.05 taken as significant.

Despite having similar bacterial numbers in the blood, mice pre-depleted of PMNs showed a significant decrease in bacterial numbers in the heart (Fig 4B). We then measured the effect of PMN depletion on serum troponin levels. As expected, there was a significant increase in the serum troponin levels in the isotype-treated, TIGR4 infected mice, compared to the uninfected control group (Fig 4C). PMN-depleted infected mice had lower troponin levels compared to their isotype-treated counterparts (Fig 4C), indicating lower cardiomyocyte damage in the animals in absence of neutrophils. Collectively, these findings point to a detrimental role for PMNs in the host cardiac tissue during invasive pneumococcal infection.

### 2.4 Late stage of invasive pneumococcal infection is associated with reduced CD73 expression on PMNs

Previous work demonstrated the importance of PMNs in controlling pulmonary bacterial numbers in the early phases of pneumococcal lung infection [10]. However, the above data with inability of PMNs to control bacterial burden in the blood (Fig 4A) and increased cardiac bacterial CFU (Fig 4B) and cardiomyocyte damage (Fig 4C) in presence of PMNs pointed to a dysregulated neutrophilic response during invasive pneumococcal infection. To test this, we examined if systemic infection with *S. pneumoniae* TIGR4 was associated with altered expression of extracellular (EAD)-pathway components on the PMNs migrating into the heart during the course of infection. The EAD-pathway plays an important role in regulating PMN responses [16] and host resistance to pneumococcal lung infection [10, 13, 21]. Using flow cytometry, we compared the expression of EAD-producing ecto-nucleotidases CD39 and CD73 on the PMNs isolated at 6-, 12- and 24hpi from the blood and heart tissue of infected animals. We found that the expression of both, CD39 and CD73 (measured by the geometric mean fluorescent intensity or gMFI), on the surface of PMNs in the cardiac tissue increased early on at 6hrs and 12hrs following infection (Fig 5A). However, as the infection progressed, we observed that the expression of these ecto-nucleotidases on the surface of PMNs was reduced by 24hpi where the levels were significantly lower compared to both uninfected baseline controls and the peak of expression following infection (Fig 5A). The decrease in CD39 and CD73 (Fig 5B) expression also occurred on circulating PMNs, indicating that the decrease in expression observed was not mediated by the tissue environment.

**Fig 5.**
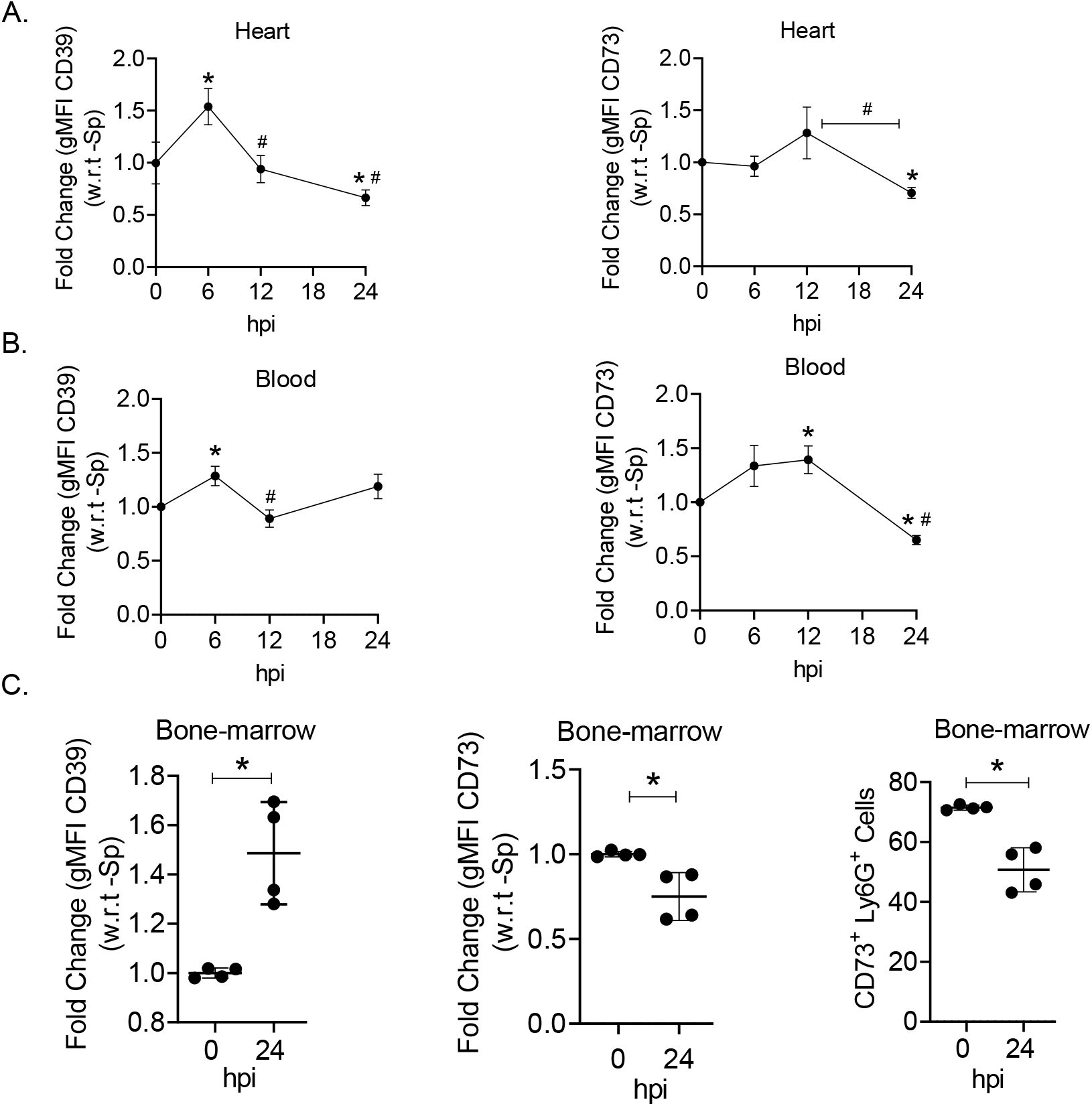
Late stage of invasive pneumococcal infection is associated with reduced expression of EAD-producing enzymes. WT (C57BL/6) mice were infected intra-peritoneally (I.P) with 10^3 CFU of *S. pneumoniae* TIGR4 or mock infected with PBS. The mice were then euthanized at 6-, 12- and 24hpi to harvest blood and heart, and at 24hpi to harvest bone-marrow. Single-cell suspension was obtained from all three tissue-types and stained with fluorescently tagged anti-Ly6G antibodies (PMNs) and antibodies against surface enzymes CD39 and CD73. Flow cytometric analysis was performed to quantify geometric mean fluorescent intensities (gMFI) of CD39 and CD73 under the Ly6G^+^ cell population in heart (A), in blood (B) and in bone-marrow (C). The line graph data in (A) and (B) are shown as Mean+/- SEM on the fold-changes of gMFI values with-respect-to the uninfected control. Data for uninfected condition are pooled from 6 mice, for 6- and 12hr are pooled from 9-13 mice, and for 24hr time point are pooled from 6-10 mice. Asterisks indicate significant difference from the baseline as calculated by One-sample t and Wilcoxon test. # indicates significant difference from the 6hr time point or the indicated group as calculated by Student’s t-test. Data in (C) are shown as Mean +/-S. D and pooled from n=4 mice per group. Asterisks indicate significant difference from the baseline as calculated by One-sample t and Wilcoxon test. *p*<0.05 taken as significant.

We next tested if the observed decrease of CD39 and CD73 on PMNs happens after PMNs have migrated from the bone-marrow or prior to their egress. We observed that systemic infection with *S. pneumoniae* TIGR4 resulted in a significant reduction in the expression of CD73 on the bone-marrow PMNs (Ly6G^+^ cells) (Fig 5C) with a significant reduction in the overall CD73^+^ PMN population in the bone-marrow (Fig 5C) by 24hrs post challenge. Interestingly, we observed a significantly higher expression of CD39 on the PMNs in the bone-marrow of infected mice (Fig 5C) suggesting that the reduction in CD39 in response to invasive pneumococcal infection occurs once the PMNs efflux into the circulation.

Collectively, these findings indicate that the later stages of invasive *S. pneumoniae* infection is accompanied by changes in the expression of EAD-pathway components on the host PMNs and that the reduction in CD73 expression on the PMNs occurs in the bone-marrow environment.

### 2.5 CD73 is required for host resistance to invasive pneumococcal infection

To test the relevance of CD73 in host resistance to invasive pneumococcal infection, we compared the course of disease in WT versus CD73^-/-^ [10], mice following intra-peritoneal challenge with 10^3 CFU of *S. pneumoniae* TIGR4. When determining the bacterial burden in the blood, we observed the presence of bacteria in only approximately 30% of WT mice at 6hpi (Fig 6A). In contrast, 80% of CD73^-/-^ mice became bacteremic within 6 hrs of infection with approximately 8-fold higher bacterial burden compared to the WT counterparts (Fig 6A). This trend was also reflected at 12hr time point where CD73^-/-^ mice had significantly higher bacterial burden in the blood compared to WT mice (Fig 6B) indicating the failure of the host to contain infection. This difference in bacterial numbers in the blood was also reflected in the overall bacterial burden in the heart, with CD73^-/-^ mice having approximately 10-fold higher CFU counts compared to WT mice (Fig 6C). However, by 24hpi we did not see any significant difference in bacterial burden between the two mouse strains in the blood (Fig 6B) and heart (Fig 6C) indicating a reduction in the capacity of WT mice to contain bacterial numbers at this later time. By this later timepoint (24hpi), CD73 expression is significantly decreased on the surface of PMNs in WT mice (Fig 5).

**Fig 6.**
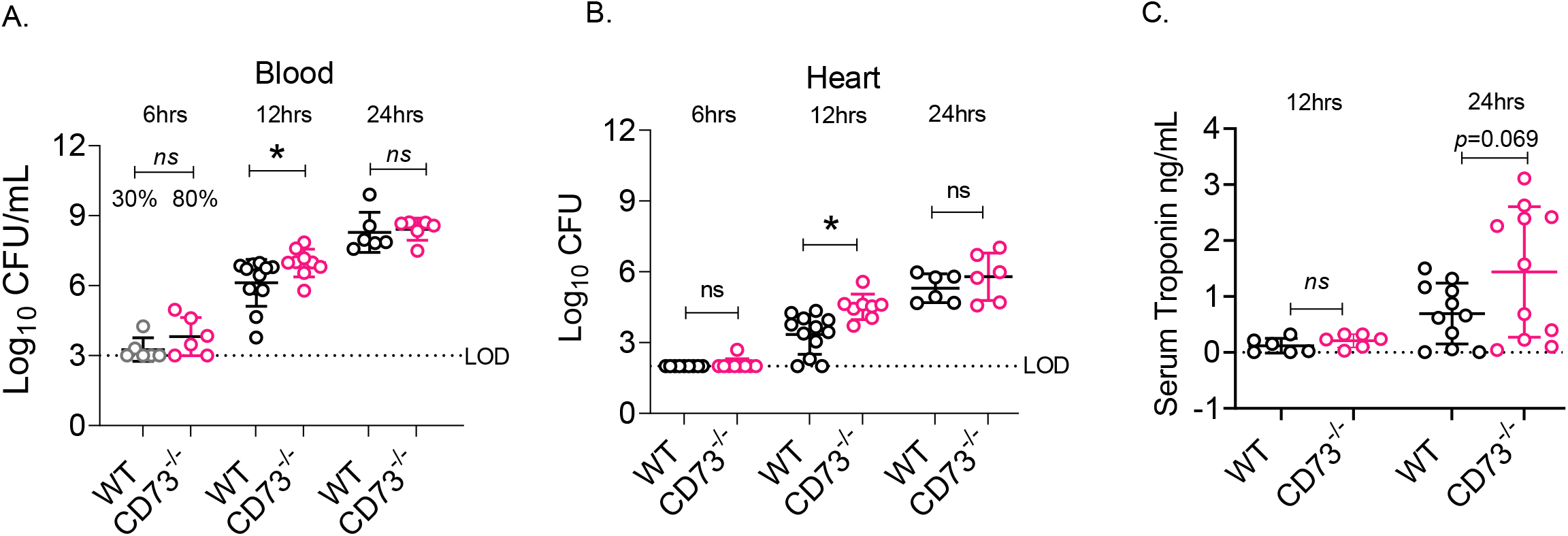
Mice lacking CD73 are more susceptible to invasive *S. pneumoniae* infection and increased cardiomyocyte damage. C57BL/6 WT and CD73^*-/-*^ mice were infected intra-peritoneally (I.P) with 10^3 CFU of *S. pneumoniae* TIGR4. The mice were then euthanized at 6-, 12- and 24hpi to harvest blood and heart to quantify **(A)** bacterial burden in the blood and **(B)** in the heart, and to quantify **(C)** the levels of Troponin-I in the sera samples. Data shown are from a total of 6-12 mice per group where every circle indicates one mouse. Percentages above the data in (A) reflect the fraction of bacteremic mice. Asterisks indicate significant differences as calculated by Student’s t-test with Welch’s correction with *p*<0.05 taken as significant.

The difference in bacterial numbers observed between WT and mice lacking CD73 was not the result of difference in the PMN migration into the cardiac tissue between the two strains as we did not observe any difference in the percentage (Fig S5) or number of PMNs in the cardiac tissues at any of the time points tested. When comparing the extent of cardiac injury, we found that CD73^-/-^ mice showed approximately 2-fold higher troponin levels (*p*= 0.07) at 24hpi compared to WT mice (Fig 6D) indicating an increased cardiac damage in mice lacking the ability to produce extracellular adenosine. Overall, these data indicate that CD73 is required to limit pneumococcal numbers and subsequent cardiac damage during invasive pneumococcal infection.

### 2.6 Late stage of invasive pneumococcal infection is associated with significantly higher levels of activated PMNs in the host cardiac tissue

PMNs can kill *S. pneumoniae* through various anti-microbial effector functions including phagocytosis, NETosis, degranulation and the release of reactive oxygen species (ROS) [22, 23]. However, these functions can also contribute significantly to the host tissue pathology [24]. We therefore, wanted to test if the influx of PMNs in the cardiac tissue during an active infection is associated with increased neutrophil degranulation which could contribute to cardiomyocyte damage. Hearts of C57BL/6 mice infected with 10^3 CFU of *S. pneumoniae* TIGR4 strain were harvested at 6-, 12- and 24hpi, and either homogenized for ELISA assays or digested to obtain single-cell population to quantify the ROS levels in the cardiac PMNs through flow cytometry. As the EAD-pathway is a known regulator of PMN functions [10, 16], the levels of anti-microbials tested were also compared to that of CD73^-/-^ mice to determine the role of CD73 in PMN responses.

When comparing the levels of neutrophil elastase (NE), a serine-proteinase stored in the azurophilic granules of PMNs, we did not observe any increase at 6hpi compared to the uninfected baseline in WT mice (Fig 7A). However, as the disease progressed, there was significant increase in NE levels at 12- and 24hr time points compared to both, uninfected and the 6hr time point (Fig 7A). Although the PMN numbers did not show an increase post 12hpi (Fig 1C), there was an approximate 2-fold increase in NE levels at 24hpi compared to 12hpi (Fig 7A). Despite having similar level of PMN presence in the hearts (Fig S5), CD73^-/-^ mice had significantly higher levels of NE at 12hrs compared to WT mice (Fig 7A) which is worth noting given the tissue damaging capacity of this protease [25]. No difference in NE levels was seen between the two mice strains by 24hpi (Fig 7A).

**Fig 7.**
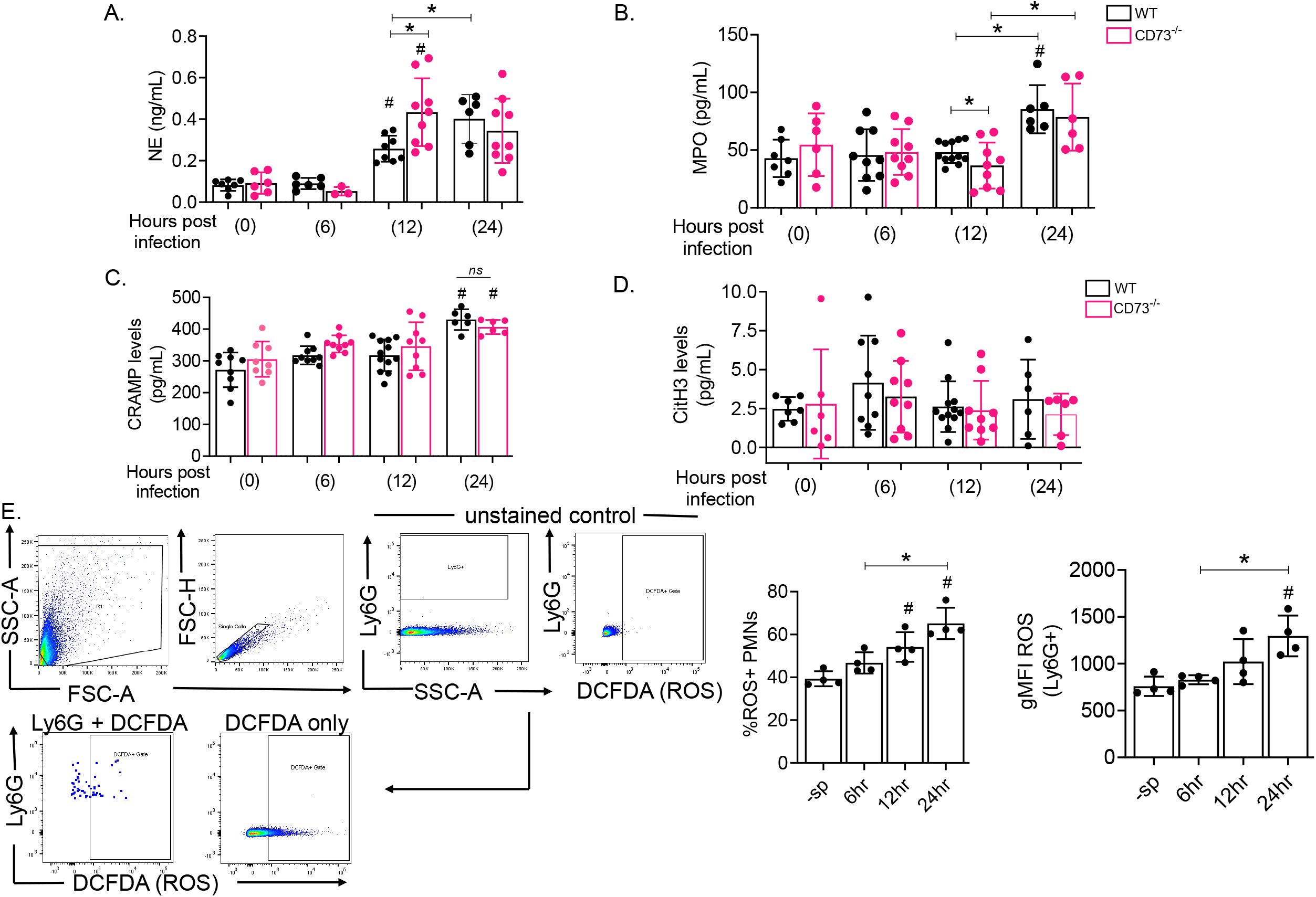
Late stage of invasive pneumococcal infection is associated with an activated PMN response. WT (C57BL/6) and CD73^-/-^ mice were infected intra-peritoneally (I.P) with 10^3 CFU of *S. pneumoniae* TIGR4 or mock infected with PBS. The mice were then euthanized at 6-, 12- and 24hpi. The hearts were to harvested and either homogenized to estimate the levels of **(A)** Neutrophil elastase (NE), **(B)** Myeloperoxidase (MPO), **(C)** Cathelicidin (CRAMP) and **(D)** Citrullinated H3 (NETosis marker) or **(E)** digested in collagenase containing buffer to obtain single cell suspension and stain with DCFDA dye (2’,7’ –dichlorofluorescin diacetate) to measure total ROS in Ly6G^+^ population using flow cytometry. Flow cytometric analysis was performed to quantify number of ROS^+^ PMNs (per the shown gating strategy) and geometric mean fluorescent intensities (gMFI) of ROS were calculated under Ly6G^+^ cell population. The data shown (A-D) are pooled from 6-10 mice per group where each circle indicate an individual mouse. The data in part E are pooled from 4 mice per time point. Multigroup analysis was done by One Way ANOVA followed by Tukey’s test while comparison between indicated two groups was done by Student’s t-test with *p*<0.05 taken as significant. * indicates significance between indicated groups and # indicates significance from the baseline.

Myeloperoxidase (MPO) is another azurophilic granule component produced by neutrophils that catalyzes the formation of a number of reactive oxidant species and plays an important role in host defense and tissue damage. We found that although PMN numbers in the cardiac tissue increased during the course of infection (Fig 1C), we did not see an increase in the MPO levels at either 6- or 12hpi compared to the uninfected baseline (Fig 7B). However, we did observe slight yet significantly lower MPO levels in CD73^-/-^ mice at 12hrs, the time point at which these mice showed higher pneumococcal burden in the heart compared to WT mice (Fig 6C). At 24hrs, the MPO levels significantly increased for both mice type compared to 12hrs with no difference between the two mouse strains (Fig 7B). The host defense peptide Cathelicidin (CRAMP) showed a similar trend as MPO, with persistent levels of Cathelicidin seen at 0-, 6- and 12hpi and a significant increase seen at 24hpi (Fig 7C), with levels similar in both mouse strains (Fig 7C).

The levels of citrullinated histoneH3, which is indicative of NETosis, were persistent at all timepoints tested (Fig 7D). However, we did not observe a significant change in citrullinated histoneH3 upon infection or between the WT versus CD73^-/-^ mice (Fig 7D).

Cardiomyocyte pathology in acute myocardial infarction is linked to increased levels of ROS in the cardiac tissue [26]. To test if there is also an increased ROS production by the PMNs migrating into the cardiac tissue, heart tissue harvested at 6-, 12- and 24hpi from WT mice were digested in collagenase solution to obtain single-cell suspension and cells were treated with DCFDA dye (2’,7’ –dichlorofluorescin diacetate) to measure total ROS by Ly6G^+^ cell population through flow cytometry. Interestingly, we saw a significant increase in the %ROS^+^ PMNs at 12- and 24hpi with significant increase in ROS levels during the late stage of invasive pneumococcal infection which coincided with the reduction in CD73 levels (Fig 5B).

Collectively, these data show that migration of PMNs in the cardiac tissue is associated with the release of various anti-microbial peptides with known role in host tissue inflammation. Additionally, significant increase in the levels of some of the tested anti-microbials was typically seen at 24hpi, the point when PMNs showed significant reduction in CD73 expression. Our data also indicate that the observed increase in some of these PMN factors at 24hpi was not merely a result of increased neutrophil numbers in the heart given that the peak PMN influx at 12hpi was not associated with increased levels of anti-microbials. Lastly, CD73^-/-^ mice having significantly altered levels of NE and MPO compared to the WT mice despite similar PMN numbers point to a possibility of EAD-mediated regulation of neutrophil function during invasive *S. pneumoniae* infection.

### 2.6 Cardiomyocytes infected with *S. pneumoniae* show higher damage in the presence of CD73^-/-^ PMNs compared to the WT PMNs

Since CD73^-/-^ mice have a systemic deletion and lack CD73 on cell types other than PMNs, we wanted to directly test the effect of CD73 expression on PMNs in cardiomyocyte damage during infection. To do that, we used the HL-1 cardiomyocyte cell line as an *ex vivo* model to study the role of PMNs and CD73 in cardiomyocyte damage. HL-1 cells have previously been used to study *S. pneumoniae*-mediated cardiomyocyte killing [19]. HL-1 cells were challenged with *S. pneumoniae* TIGR4 strain at a multiplicity of infection (MOI) of 10 for 10 minutes with and without the presence of bone-marrow derived PMNs supplemented with 3% sera (to opsonize pneumococci), to closely mimic *in vivo* conditions. Following incubation, the HL-1 cells were washed, trypsinized and stained with Annexin-PI to enumerate the cellular viability (Fig 8A). PMNs from CD73^-/-^ were also used in these assays to determine the role of CD73 in regulating PMN response and subsequent cardiomyocyte damage.

**Fig 8.**
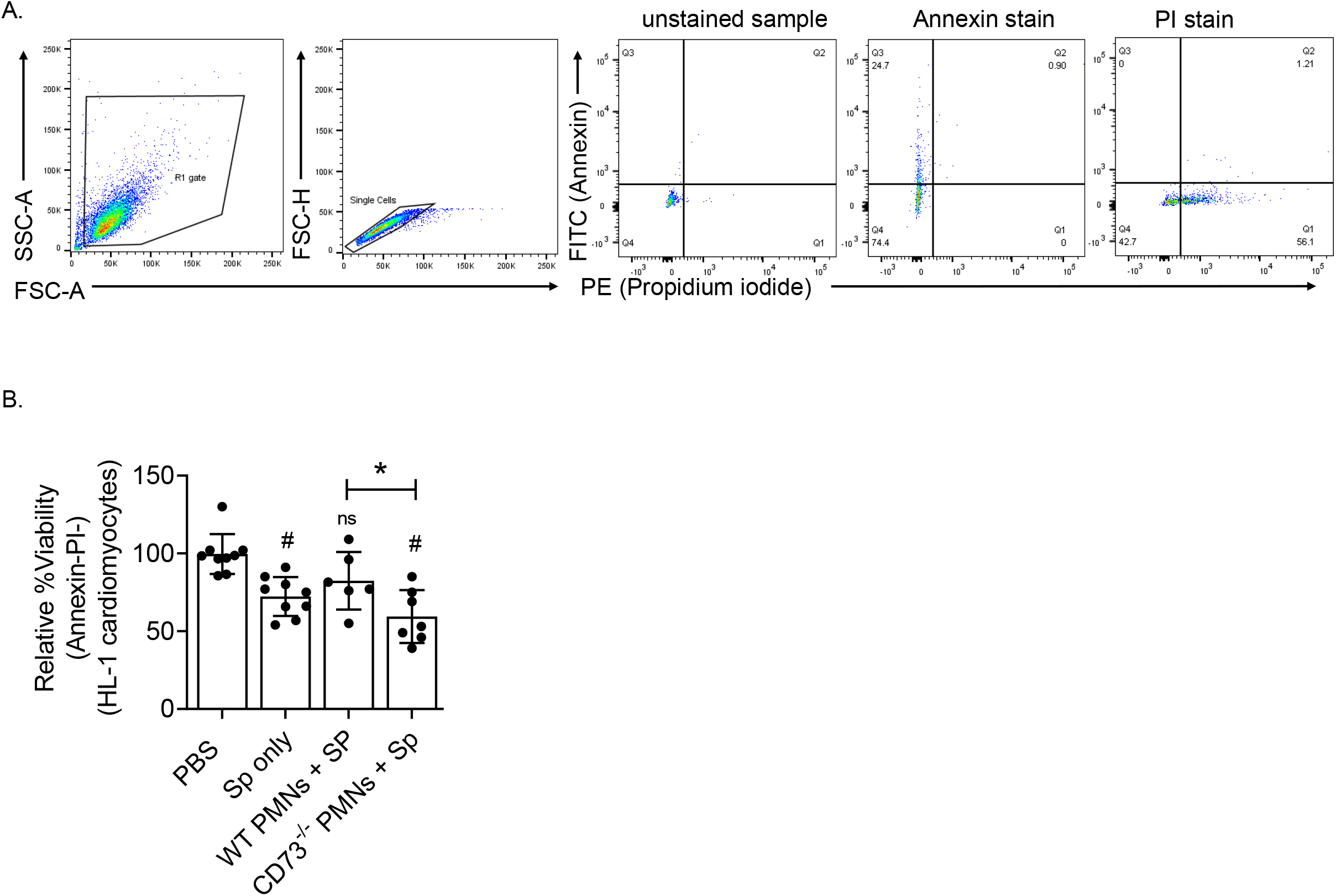
PMNs from CD73^-/-^ mice exacerbate pneumococci-induced cardiomyocyte cell death. PMNs isolated from the bone marrow of the indicated strains of mice were incubated for 10 min at 37°C with HL-1 cardiomyocytes infected with *S. pneumoniae* TIGR4 (in presence of 3% mouse serum) at an MOI of 10 or were mock treated (uninfected) with 3% matching mouse serum only. Flow cytometry was used to determine the effect of PMN presence on HL-1 cardiomyocyte cell viability. HL-1 cells were washed extensively to remove PMNs before trypsinizing and staining with Annexin-PI stain (A) Gating strategy followed during analysis of flow cytometry data where the % of Live cells in Q4 (Annexin-PI-) were quantified and compared between groups (B). Data shown are pooled from three biologically independent experiments where each condition was tested at least in duplicate. The percent of live cells for each condition was normalized to the PBS condition within the same experiment. * indicates significance between indicated groups and # indicates significance from the PBS group as calculated by One Way ANOVA followed by Tukey’s test. *p*<0.05 taken as significant.

Infection of HL-1 cells with *S. pneumoniae* alone caused a significant cellular damage resulting in reduction in cardiomyocyte viability compared to the PBS treated control (Fig 8B), as shown in previous studies [19]. Addition of PMNs from the WT mice to the infected HL-1 cells did not result in further reduction in cardiomyocyte viability compared to the cells challenged with *S. pneumoniae* alone (Fig 8B). However, when HL-1 cells were co-incubated with *S. pneumoniae* and CD73^-/-^ PMNs, there was an additional significant reduction in cardiomyocyte viability compared to the WT PMN condition (Fig 8B) indicating higher cellular damage in the presence of CD73^-/-^ PMNs. The difference in cardiomyocyte toxicity observed in presence of these two PMN types was neither due to difference in their viability (Fig S6A) nor to the number of bacteria that were able to bind to the HL-1 cells (Fig S6B). Collectively, these findings suggest that loss of CD73 expression on PMNs renders these cells damaging to cardiomyocytes during *S. pneumoniae* infection.

## 3. Discussion

The last few years have seen an increased interest in elucidating host-pathogen interactions during invasive *S. pneumoniae* infections including those involving cardiac tissue [3, 27-29]. Most of the previous work on infectious endocarditis has been focused on viridans and group A *Streptococci* and *S. aureus* which are capable of entering the blood stream of an individual either through endothelial cell injury or poor oral hygiene [30, 31]. Once in the bloodstream, these bacteria are able to attach to the inner lining of the heart directly or through abnormal heart valves and pre-existing microscopic clots [32]. The ability of *S. pneumoniae*, a leading cause of bacterial pneumonia, to infect the heart, is gaining attention given its ability to utilize the host receptors in the pulmonary environment to reach the blood-stream [9, 27, 28, 33]. Once in the blood stream, *S. pneumoniae* through pneumococcal adhesin protein (CbpA) binds to host ligands, Laminin receptor and Platelet-activating factor receptor on vascular endothelium, to infect the myocardial tissue leading to cardiomyocyte damage [7]. This is followed by pneumolysin-mediated necroptosis of macrophages allowing the bacteria to blunt the host response in the heart [8]. The majority of the studies have focused so far on bacterial factors and their effect on macrophages in the cardiac tissues, however the role of PMNs in these infections has not been fully characterized. The aim of this study was to examine the role of PMNs in *S. pneumoniae* infection of the host cardiac tissue and to elucidate the role of EAD, a host signaling pathway, in regulating PMN responses. We found that the presence of PMNs in cardiac tissue was detrimental to the host and exacerbated cardiomyocyte damage. We further found that the damaging effect of PMNs on cardiac tissue was associated with reduced expression of the EAD producing enzyme CD73 on PMNs during the later phases of disease. This is the first report, to the best of our knowledge, which shows that PMNs exacerbates pneumococci-induced cardiomyocyte damage and that the detrimental role of PMNs is due to changes on the EAD-pathway on these cells.

We found that PMNs homed to the infected cardiac tissue following invasive pneumococcal infection. PMN numbers in the cardiac tissue did not increase until 12hr of infection despite increase in the numbers of circulating PMNs early during the course of infection. This could be due to the lack of presence of *S. pneumoniae* in the cardiac tissue at the early time point when their numbers are still on the lower side in the blood. Given the different functional roles these cells play in host immunity [34, 35], the increased influx of PMNs in the cardiac tissue with the progression of disease seems to be in response to infection of myocardium by *S. pneumoniae*. Prior studies showed the presence of PMNs in the cardiac tissue of *S. pneumoniae* infected mice followed by their disappearance from the heart [8]. Here we found PMNs in the cardiac tissue at all the time points tested. With severe clinical symptoms associated with the invasive pneumococcal infection, the mice in our study were euthanized at 24hpi instead of later time points in the above-mentioned study [8]. Given that we observed persistent presence of PMN degranulation products throughout the infection, it is possible that invasive infection of *S. pneumoniae* results in multiple waves of neutrophils migrating to the heart. The late stage of infection was associated with the egress of immature PMNs with reduced surface expression of CD73, which could be a potential consequence of emergency granulopoiesis to meet the high demand of the host [36]. In an ideal situation, the influx of PMNs at the site of infection involves the control of infection by these cells through various anti-microbial functions followed by their resolution from the tissue. In situations of acute inflammation, the clearing of bacteria and cellular debris by PMNs is followed by resolution and tissue homeostasis [37, 38]. Macrophages, through the process of efferocytosis, engulf apoptotic neutrophils to activate host anti-inflammatory response [39, 40]. In invasive pneumococcal infection, once *S. pneumoniae* translocate into the myocardium, they not only damage cardiomyocytes but also kill macrophages through the release of the pore-forming toxin pneumolysin, producing cell-free zones around the cardiac microlesions [8]. This could possibly impair the regulated and coordinated clearing of PMNs from the cardiac tissue potentially leading to PMN-mediated increased cardiac inflammation giving the pathogen an upper hand over the host immune system. Further studies are required to dissect the host and bacterial factors mediating PMN influx and resolution from cardiac tissues.

We found that PMNs failed to efficiently control bacterial numbers in the cardiac tissue during systemic infection. Despite the increase in PMN numbers in the cardiac tissues, the bacterial burden increased pointing to the reduced ability of these cells to control the infection. In fact, PMN depletion did not impair bacterial clearance from the heart but rather resulted in reduced bacterial numbers suggesting a change in PMN phenotype from antimicrobial to promoting an environment more permissive to bacterial growth as the infection progressed. When looking for mechanisms, we found a significant reduction in the expression of CD73 on PMNs during the late stage of invasive pneumococcal infection. CD73 is a host surface enzyme which has been shown to be important for anti-pneumococcal functions of neutrophils in pulmonary infections [12]. Reduced expression of CD73 on the PMNs migrating into the lungs of animals infected with *S. pneumoniae* was associated with increased levels of anti-inflammatory IL-10 which significantly reduced the capacity of these cells to kill *S. pneumoniae* both, *in vivo* and *ex vivo* [12]. Additionally, pharmacological inhibition of CD73 on PMNs isolated from healthy human donors reduces their capacity to kill *S. pneumoniae* in *ex vivo* killing assays [12]. This reduced killing by the PMNs could mean more *S. pneumoniae* available to infect and damage the myocardial cells during systemic infection. In support of that, in this study, CD73^-/-^ mice had significantly higher numbers of bacteria in the heart and blood compared to WT controls within the first 12 hours of infection. This difference in bacterial control was lost by 24hrs, a timepoint that coincided with the loss of CD73 expression on PMNs in WT mice. Since adenosine can act through any of the four adenosine receptors (A1, A2A, A2B and A3) [16], future work will be aimed at finding the adenosine receptor regulating the PMN responses during invasive pneumococcal infection.

The cardiac influx of PMNs during systemic pneumococcal infection correlated with cardiomyocyte damage. This damage could be directly mediated by PMNs acting on cardiomyocytes. The act of PMN-mediated bacterial killing itself involves the release of plethora of anti-microbial peptides, proteolytic enzymes and reactive oxygen species [22]. However, in doing so, these cells are capable of causing collateral damage to the host cells [24] including cardiomyocytes. Adverse cardiac remodeling and cardiomyocyte death following reperfusion in acute myocardial infarction injury is linked to oxidative stress which in part is regulated by PMN-derived MPO [26] or aggravation of cardiac inflammation through release of pro-inflammatory mediators and extracellular vesicles by these cells [34]. Studies using either the approach of PMN depletion [41, 42] or blocking the activation of these cells [43] showed reduction in the intensity of ischemic myocardial injury in the animal models attributed to the reduction in the levels of PMN effectors in the cardiac tissue. In systemic pneumococcal infection, we saw a consistent presence of PMN degranulation products with significant increase in the levels of ROS, MPO, ROS, NE and Cathelicidin in the cardiac tissues at the 24hr time point. This time point was linked to the most increase in serum troponin levels, which is not surprising given the tissue injury capacity of the factors released by PMNs [38]. The increase in the degranulation products and tissue damage in the cardiac tissue coincided with significantly reduced CD73 levels on the PMNs in the WT mice. Further, CD73^-/-^ mice had significantly increased levels of MPO and NE in cardiac tissue and increased serum troponin levels despite similar numbers of PMNs compared to WT mice. This suggests that the absence of CD73 renders PMNs hyper activated, potentiating their ability to cause cardiomyocyte damage. This could be challenging to assess *in vivo* given the difference in bacterial burdens in the absence of CD73. Our findings *ex vivo* with HL-1 cells suggest that CD73 expression on PMNs is required for preventing cardiomyocyte damage in the context of infection and that this is independent of bacterial numbers. It is possible that CD73 is required to control PMN responses and / or alternatively EAD produced by CD73 on PMNs is signaling via adenosine receptors on cardiomyocytes and modulating their response to bacterial damage. Future work will focus on elucidating the mechanism by which EAD production and signaling controls cardiomyocyte damage during pneumococcal infection.

PMNs could alternatively control cardiomyocyte damage via their effect on bacterial numbers. Depletion of PMNs significantly reduced cardiomyocyte damage during systemic pneumococcal infection and this was associated with lower bacterial numbers in depleted hosts. *S. pneumoniae* TIGR4 form microlesions in the heart after crossing endothelium and translocating into the myocardial tissue [7, 20]. Therefore, the reduced serum troponin levels in PMN depleted mice could be due to less bacteria being able to reach and damage the host myocardium. In ischemic myocardial injury model, PMNs following activation were found to aggregate and adhere to endothelium resulting in plugging of capillary walls [44]. This raises an interesting possibility that PMN-mediated damage of the endothelium could be responsible for providing increased bacterial access to cardiomyocytes. During pulmonary infection, influx of PMNs across pulmonary endothelial and epithelial cells resulted in bacterial egress in the opposite direction promoting systemic spread of the infection [10, 45]. This could represent a novel point of intervention to prevent excessive cardiac tissue damage during invasive pneumococcal infection. Future work will be aimed at studying the role of PMNs and EAD-pathway on endothelial cell damage during *S. pneumoniae* infection.

The role of PMNs in bacterial infective endocarditis has been examined in the context of a few infections. Immunostaining on the biopsy samples from the valve of patients suffering from endocarditis due to various infectious etiologies, including *Staphylococci* and *Streptococci*, showed massive infiltration of PMNs in the areas of bacterial growth, also known as septic thrombi or vegetations [46]. These areas of vegetations were associated with high levels of elastase and MPO secreted by PMNs [46]. Interestingly, apart from controlling bacterial numbers, the serine proteases and nucleosomes released by PMNs through inactivation of coagulation suppression pathway, can lead to large-vessel thrombosis potentially increasing the risk of myocardial infarction [47]. In fact, in two separate retrospective case control studies, the infective endocarditis patients with bad clinical outcome had higher value of neutrophil-to-lymphocyte ratio in blood as compared to the ones with good clinical outcome [48, 49], indicating the risk of increased clinical severity associated with increased PMN numbers in endocarditis patients. *S. agalactiae* or Group B *Streptococcus*, a leading cause of neonatal sepsis, causes exotoxin-induced cardiomyocyte damage with poor outcome [50], a clinical feature similar to invasive *S. pneumoniae* infection. However, the role of host immune response, especially that of neutrophils in cardiomyocyte damage induced by *S. agalactiae* is not well known. This bacterium however, is known to produce C5a peptidase and CspA serine protease that impair or blunt PMN recruitment and response at the site of infection [51]. In *S. aureus*-mediated mouse endocarditis model, infiltration of PMNs in infected thrombi is shown to be important to limit the disease progression [52]. However, this influx of leukocytes around the foci of infection was also accompanied by extensive proteolytic cardiac tissue damage and cellular apoptosis [52]. On the other hand, during infective endocarditis caused by *E. faecalis*, release of metalloproteinases and elastase by PMNs infiltrating in the cardiac tissue were suggested to promote the formation and growth of bacteria containing thrombi in the heart, and induce myocardial damage [53]. These previous studies point towards a detrimental role of neutrophils in the cardiac tissue, which is supported by our findings here in the context of *S. pneumoniae* infection. However, the effect of PMN migration on the bacterial burden per se seems to be dependent on the virulence of the causative agent involved in the infection.

In summary, this study demonstrated the detrimental role of PMNs in host cardiac tissue during invasive pneumococcal infection causing cardiomyocyte damage. Importantly, we identified CD73 as a regulator of this damaging response and a potential target to therapeutically influence PMN function to efficiently manage tissue damage during invasive pneumococcal infection.

## 4. Materials and Methods

### 4.1 Mice

All the mice work was conducted in accordance per the guidelines set by Institutional Animal Care and Use Committee (IACUC), approval number MIC33018Y. Wild-type (WT) C57BL/6 (B6) mice were purchased from Jackson Laboratories (Bar Harbor, ME, USA). CD73^-/-^ mice on C57BL/6 background [54] were purchased from Jackson Laboratories and bred at a specific-pathogen free facility at the University at Buffalo. Both, male and female mice of 8-12 weeks of age were used in this study.

### 4.2 Bacteria

*Streptococcus pneumoniae* serotypes 4 (TIGR4) and 2 (D39) were used in this study [55]. Bacteria were grown in Todd-Hewitt broth (BD Biosciences) supplemented with Oxyrase and 0.5% yeast extract at 37°C/5% CO_2_. Once in the mid-log phase, bacterial aliquots were frozen at –80°C in growth media with 20% glycerol. Prior use, the aliquots were thawed, washed and diluted in phosphate buffer saline (PBS) to reach desired CFU and the numbers were confirmed by plating the bacterial suspension on tryptic soy agar plates with 5% sheep blood (Hardy Diagnostics).

### 4.3 Infection

Mice were challenged intraperitoneally (i.p) with 10^3^ CFU of mid-log phase grown *S. pneumoniae* in 100 μl of PBS to establish invasive pneumococcal infection and recapitulate the cardiac damage in a synchronized manner, as previously described [9, 20]. Mice were euthanized at 6-, 12- and 24hr-post infection (hpi) to collect blood, bone marrow and heart tissue. Prior to collecting heart tissue, mice were perfused with sterile PBS by inserting the needle in the left ventricle and cutting the right atrium to ensure the removal of blood-circulating bacteria from coronary vasculature, as previously described [20]. For lung infection, mice were intratracheally (i.t.) challenged with *S. pneumoniae* TIGR4 at dose of 1 × 10^5^ CFU [10] and euthanized at 24hpi to detect the presence of PMNs in the heart tissue through Flow cytometry.

### 4.4 Flow cytometry

Flow cytometry was used to determine the numbers of PMNs, the expression of EAD-pathway components on the PMNs and cardiomyocytes, and measurement of total ROS produced by the PMNs isolated from the blood and heart of infected animals euthanized at 6-, 12- and 24hpi. Blood (300μl per mouse) was first collected by portal vein snips using 50mM EDTA/PBS as an anticoagulant [56] followed by the perfusion and harvesting of the heart tissue, as explained above. The hearts were longitudinally cut and one half homogenized to determine bacterial numbers [19] while the other half minced into pieces and digested for 20 minutes with RPMI 1640 media with 10% FBS, 1 mg/ml Type II collagenase (Worthington), and 50 U/ml Deoxyribonuclease I (Worthington) at 37°C/ 5% CO2 to get a single cell suspension. Red blood cells from the collected blood or from the single cell suspension of the cardiac tissue were removed via treatment with a hypotonic lysis buffer (Lonza). Cells were then washed and the following antibodies were used for flow cytometry; Fc block (anti-mouse clone 2.4G2), Ly6G (clone IA8; invitrogen), CD73 (clone TY/11.8; eBioscience) and CD39 (clone24DMS-1; eBioscience). Fluorescence intensities were measured on a BD FACS Fortessa and data were analyzed using FlowJo. At least 10k events were analyzed using FlowJo for all sample types. For experiments requiring PMN depletion, mice were treated intraperitoneally with 50 µg of the Ly6G-depleting antibody IA8 or isotype IgG control (BioXCell) [57], a day prior and at the time of infection with *S. pneumoniae* and PMN depletion was confirmed using anti-Gr-1 Rb6-antibodies (Invitrogen).

### 4.5 Measurement of PMN effector functions

Heart homogenates were aliquoted and stored at -80°C. Aliquots were thawed on ice, centrifuged briefly at 3000rpm for 2 minutes to settle the tissue debris. The supernatants were then collected and used to determine Cathelicidin antimicrobial peptide (CRAMP) (CusaBio), myeloperoxidase (MPO) (invitrogen), citrullinated histone H3 (citH3) (Cayman) and neutrophil elastase (NE) (R&D Systems) levels using the commercially available ELISA quantification kits using the manufacturer’s instructions. For examining total ROS production by PMNs *in vivo*, mice were euthanized at 6-, 12- and 24hpi following invasive pneumococcal pneumonia. The heart tissue were then minced into pieces and digested for 20 minutes with RPMI 1640 supplemented with 10% FBS, 1 mg/mL Type II collagenase (Worthington), and 50 U/mL Deoxyribonuclease I (Worthington) at 37°C/ 5% CO2, as previously described [56]. Single-cell suspensions were resuspended in FACS buffer (HBSS/ 1% FBS), treated with Fc block and surface stained with for Ly6G and DCFDA dye (2’,7’ –dichlorofluorescin diacetate) (total ROS) for 30 minutes at 37°C. Cells were then washed and resuspended in FACS buffer and ROS production measured using flow cytometry.

### 4.6 PMN isolation

For all the *ex vivo* experiments involving HL-1 cells and *S. pneumoniae* TIGR4, PMNs were isolated from the bone marrow of WT and CD73^-/-^ mice, as previously described [13, 58]. Briefly, the cell suspension obtained from the bone marrow of mice was treated with hypotonic lysis buffer to get rid of red blood cells. The remaining cell suspension was then layered on Histopaque 1119 and Histopaque 1077 and PMNs were isolated through density gradient centrifugation and re-suspended in Hanks’ Balanced Salt Solution (HBSS)/0.1% gelatin without Ca^2+^ and Mg^2+^, and used in subsequent assays. This method yields 85-90% of enriched PMN population as measured by flow cytometry using Ly6G and CD11b antibodies [12].

### 4.7 HL-1 Cells

PMN-mediated toxicity of cardiomyocytes in presence of *S. pneumoniae* was determined using mouse atrial cardiomyocytes (HL-1) cells (Millipore Sigma). HL-1 cells HL-1 cells were passaged and grown in Claycomb media (Sigma) supplemented with 10% of HL-1 qualified FBS (EMD Millipore), norepinephrine (Sigma), L-glutamine (EMD Millipore) and Penicillin/Streptomycin (EMD Millipore) as per manufacturer’s instructions. HL-1 cells were grown on gelatin/fibronectin coated surface of either T25 flask (passaging) or 96-well plate (*ex vivo* assay). Trypsinization of HL-1 cells was done using HL-1 qualified trypsin/EDTA solution (Sigma).

### 4.8 Cytotoxicity assay using HL-1 cells

Cytotoxicity assay using HL-1 cells was done per previously defined protocol [59] with slight modification. Briefly, HL-1 cells were seeded in a 96-well plate at a cell density of 1.5 × 10^5^ cells/well, a day prior to the experiment. The cells were incubated overnight at 37°C with 5% CO_2_ to allow adherence to the plate surface. The following day, the cells were washed twice with PBS and challenged with 2.5 × 10^5^ of bone-marrow derived PMNs [58, 60] with or without the presence of *S. pneumoniae* TIGR4 strain (MOI of 4) in plain Claycomb media (without FBS and antibiotics) for 10 minutes at 37°C with 5% CO_2_. The cells were then extensively washed, trypsinized and stained with Annexin-PI staining (BioLegend) to determine cell viability.

### 4.9 Bacterial binding assay in presence of PMNs

Standard bacterial binding assay [61] was used with slight modifications to enumerate the binding of bacteria to HL-1 cells in presence of PMNs. Briefly, HL-1 cells were seeded in a 96-well plate at a cell density of 1.5 × 10^5^ cells/well, a day prior to the experiment and allowed to adhere to the plate surface overnight at 37°C with 5% CO_2_. The following day, the cells were washed twice with PBS and challenged with 1 × 10^5^ of bone-marrow derived PMNs and 1 × 10^3^ CFU of *S. pneumoniae* (pre-opsonized win 3% mouse sera) in 100μl of plain Claycomb media (without FBS and antibiotics). The plates were incubated for 45 minutes at 37°C. To determine the number of bacteria that adhered to cardiomyocytes, the supernatants were discarded and cells were washed five times with PBS and lifted with HL-1 qualified trypsin/EDTA solution. The cell volumes were vigorously pipetted to produce a homogenous suspension which were then serially diluted and plated on blood agar plates. The percent of bacteria bound to HL-1 cells were determined with respect to no cell control well where bacteria were incubated under the same experimental conditions and enumerated on blood agar.

### 4.10 Measurement of cardiac damage

Troponin-I (cTn-I) levels (Life Diagnostics), as a measure of cardiac damage [8, 9], were measured using the sera collected from mice euthanized at 6-, 12- and 24hpi with *S. pneumoniae* and stored at -80°C. The ELISA for serum Troponin-I was run per the manufacturer’s protocol.

### 4.11 Statistical Analysis

Graph Pad Prism v. 9 was used to perform all the statistical analysis. The data shown are pooled from separate experiments, unless specified. The data in the line graphs, bar graphs and dot plots are shown as mean values +/- standard deviation (SD) unless specified. Bacterial numbers were log-transformed. Distribution of the data was tested using the D’Agostino & Pearson and Shapiro-Wilk tests. Significant difference between two groups was determined by unpaired parametric Student’s t-test while multi-group comparison was done with one-way ANOVA followed by Tukey’s multiple comparison test as indicated. All the correlation analysis was done using Pearson correlation analysis. *P* values < 0.05 were considered significant. Each experiment was done at least three independent times. The number of biological and technical replicates per experiment is indicated in each figure legend.

## 5. Acknowledgements

We would like to acknowledge Shaunna Simmons and Sydney Herring for their critical feedback on the manuscript.

## 7. Author contributions

MB designed and conducted research, analyzed data and wrote paper. VRR conducted research, analyzed data and wrote paper. AL and EYIT conducted research and analyzed data. JL provided valuable reagents, helped with the data interpretation, and provided feedback. ENBG designed research and wrote the paper. MB had responsibility for final content. All authors read and approved the final manuscript.

