## Supplementary material for "Neutrophil-driven cardiac damage during invasive *Streptococcus pneumoniae* infection is regulated by CD73": Paper Figures 12.5.22 Supplementary.pptx

### Slide 1
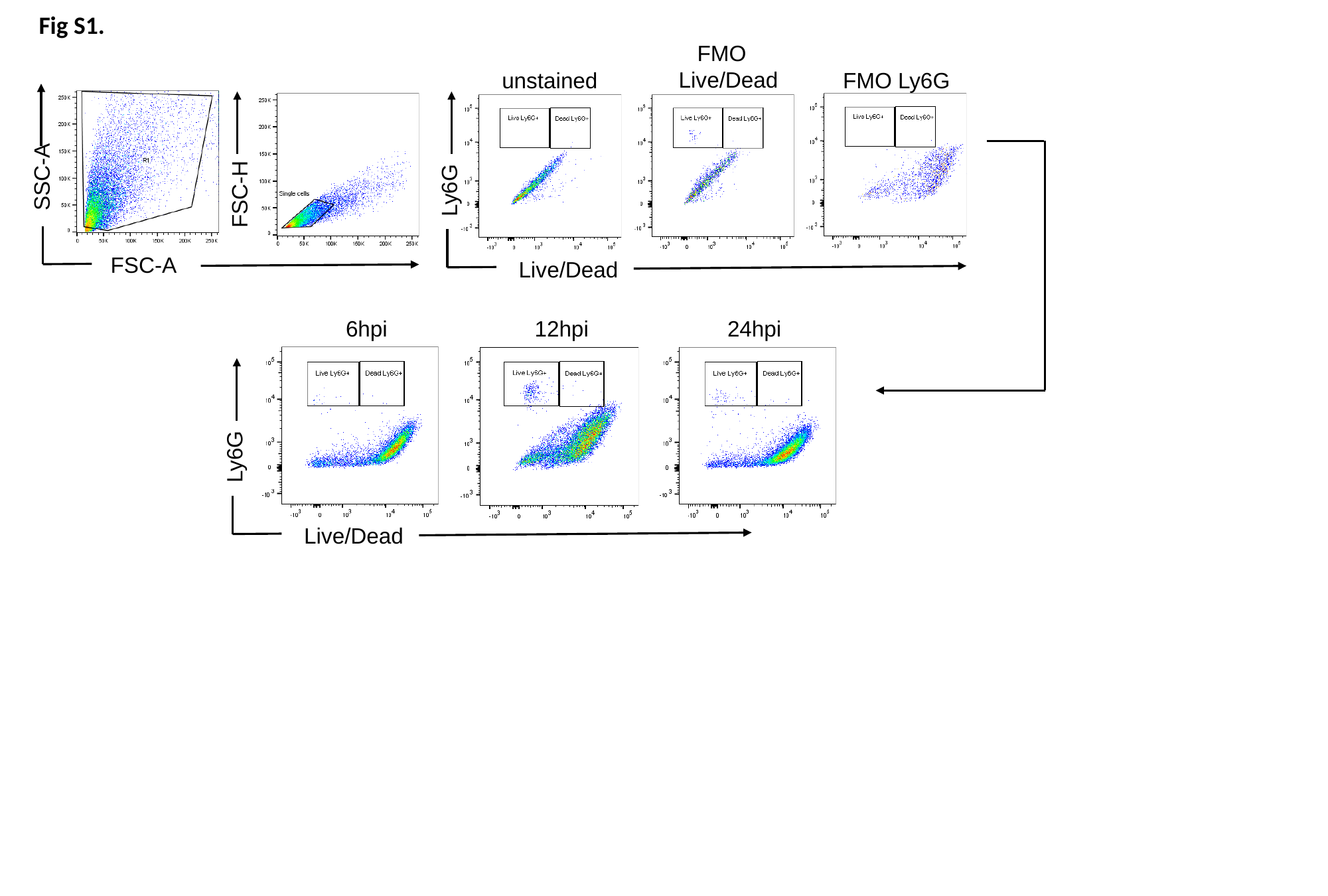

Fig S1.
 FMO
Live/Dead
unstained
 FMO Ly6G
SSC-A
Ly6G
FSC-H
FSC-A
Live/Dead
6hpi
12hpi
24hpi
Ly6G
Live/Dead

### Slide 2
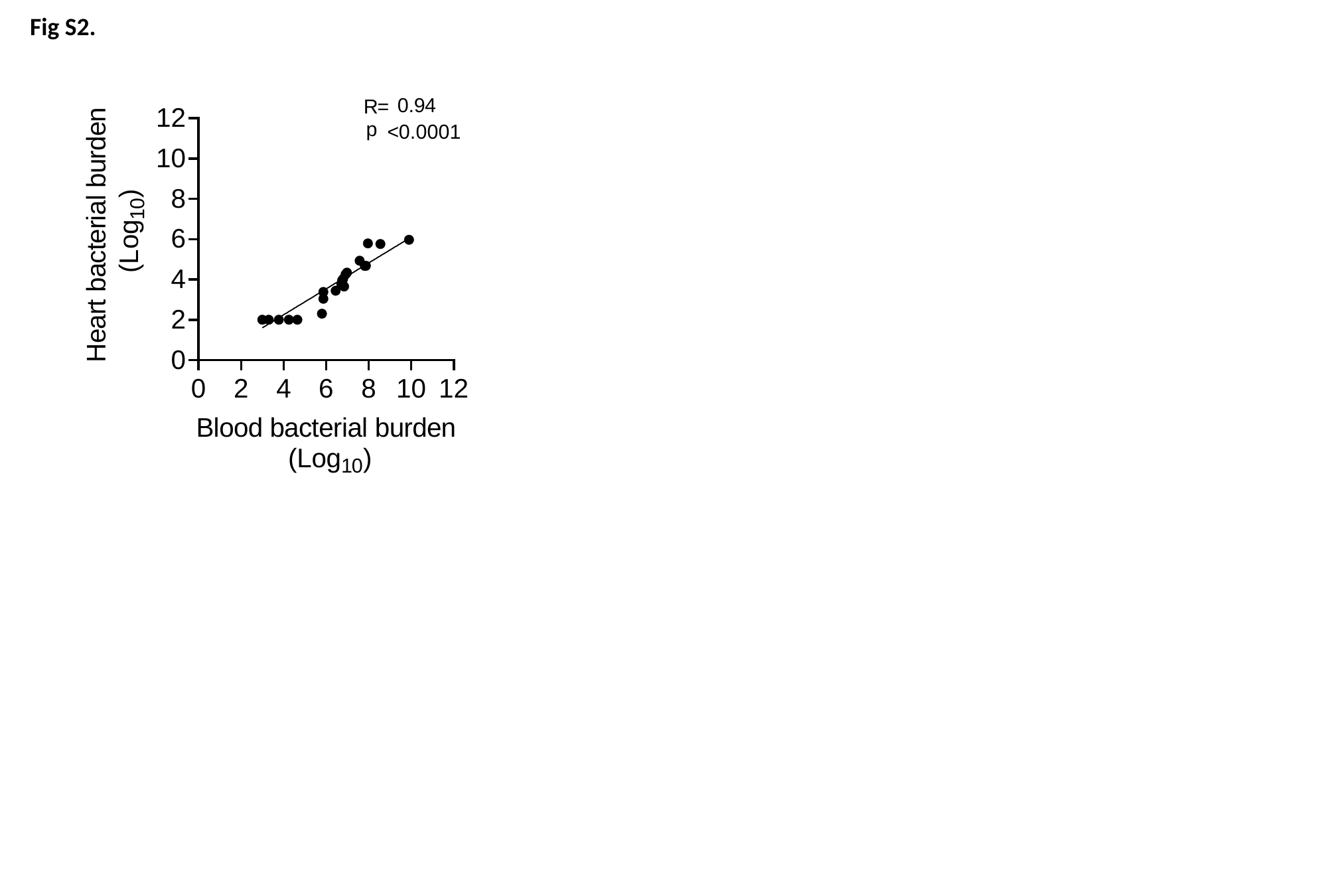

Fig S2.

### Slide 3
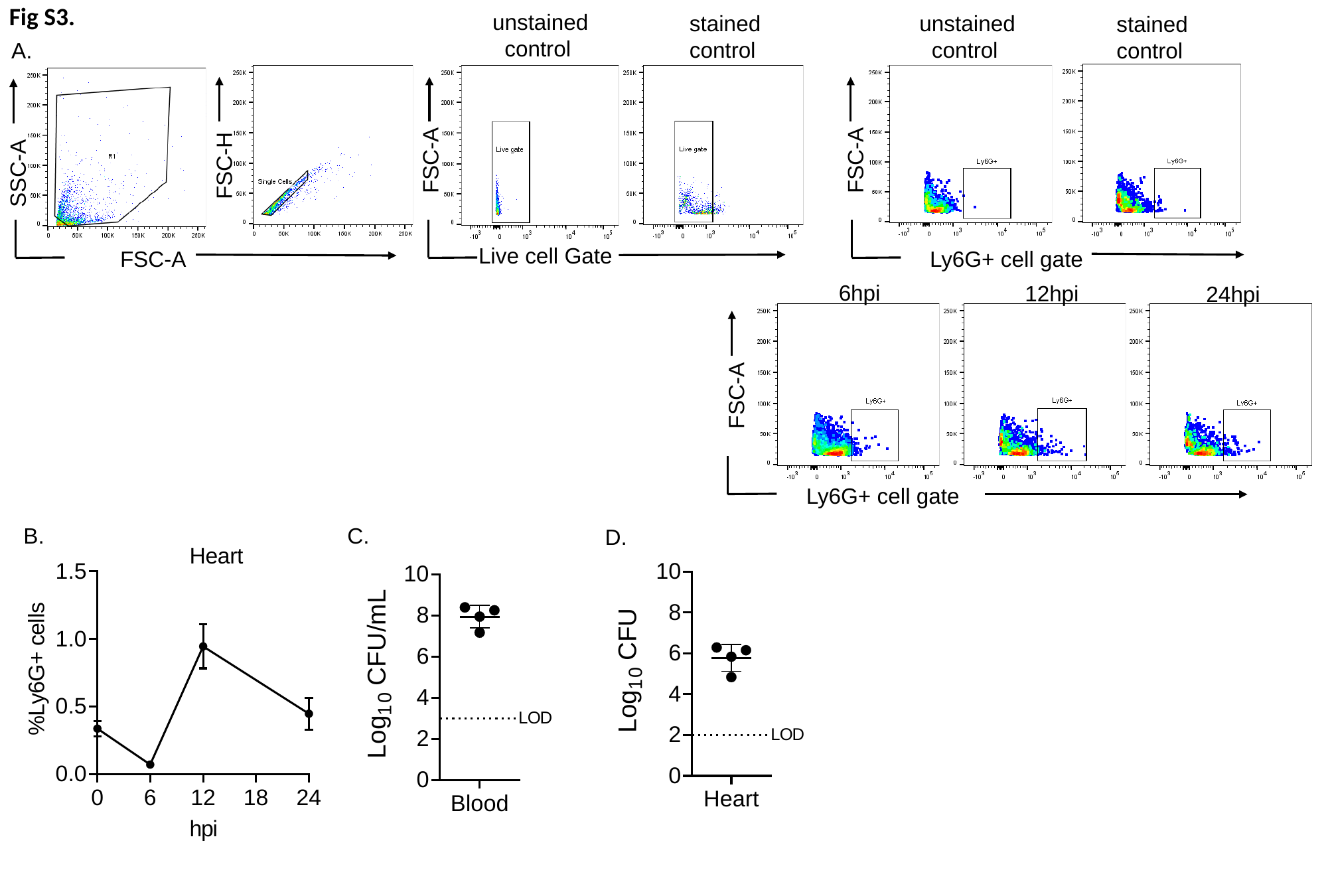

Fig S3.
unstained
 control
unstained
 control
stained
control
stained
control
A.
FSC-A
FSC-A
FSC-H
SSC-A
Live cell Gate
FSC-A
Ly6G+ cell gate
6hpi
12hpi
24hpi
FSC-A
Ly6G+ cell gate
B.
C.
D.

### Slide 4
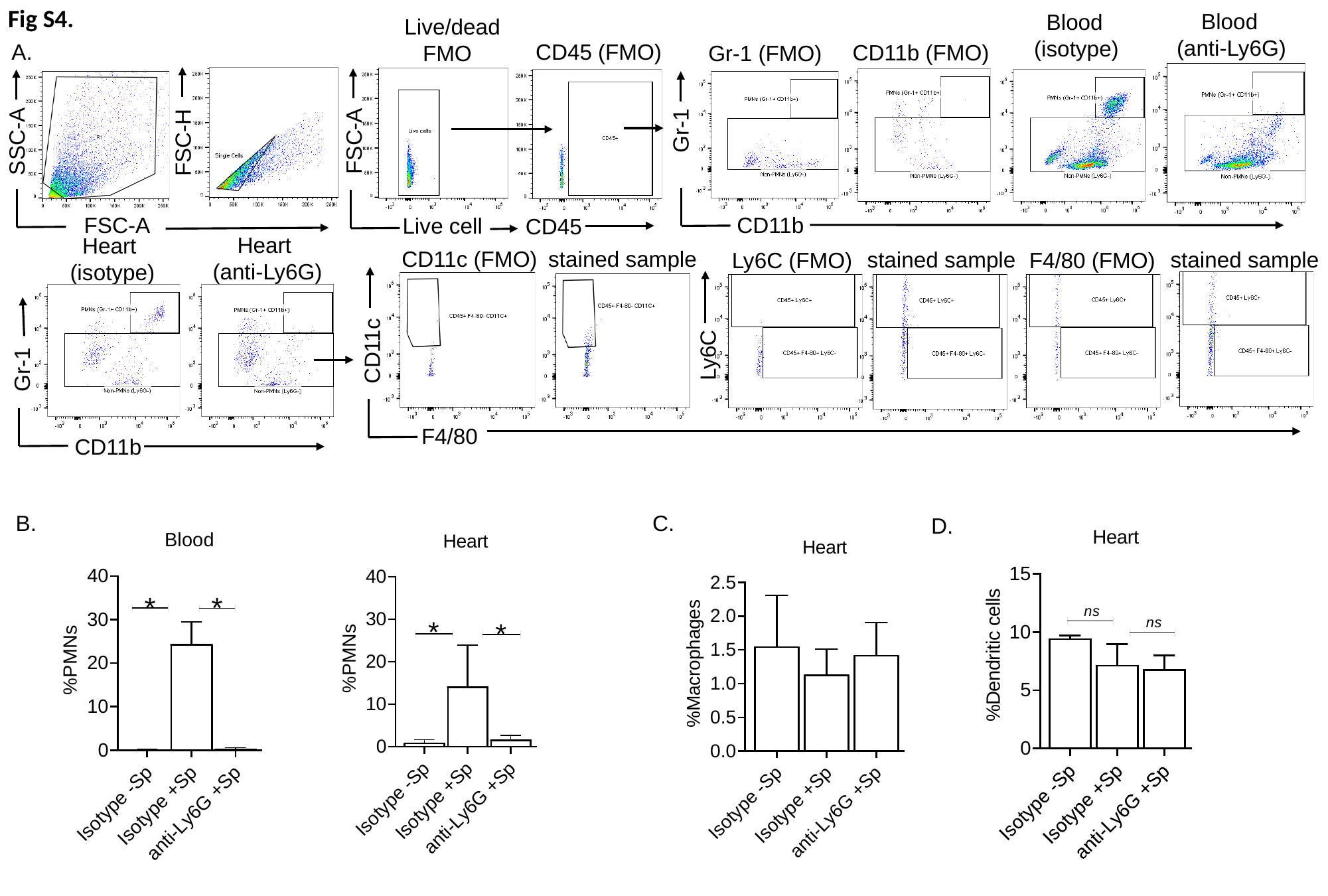

Fig S4.
 Blood
(anti-Ly6G)
 Blood
(isotype)
Live/dead
 FMO
A.
CD45 (FMO)
CD11b (FMO)
Gr-1 (FMO)
Gr-1
FSC-A
SSC-A
FSC-H
CD11b
FSC-A
Live cell
CD45
 Heart
(anti-Ly6G)
 Heart
(isotype)
Gr-1
CD11b
stained sample
CD11c (FMO)
stained sample
stained sample
Ly6C (FMO)
F4/80 (FMO)
CD11c
Ly6C
F4/80
B.
C.
D.

### Slide 5
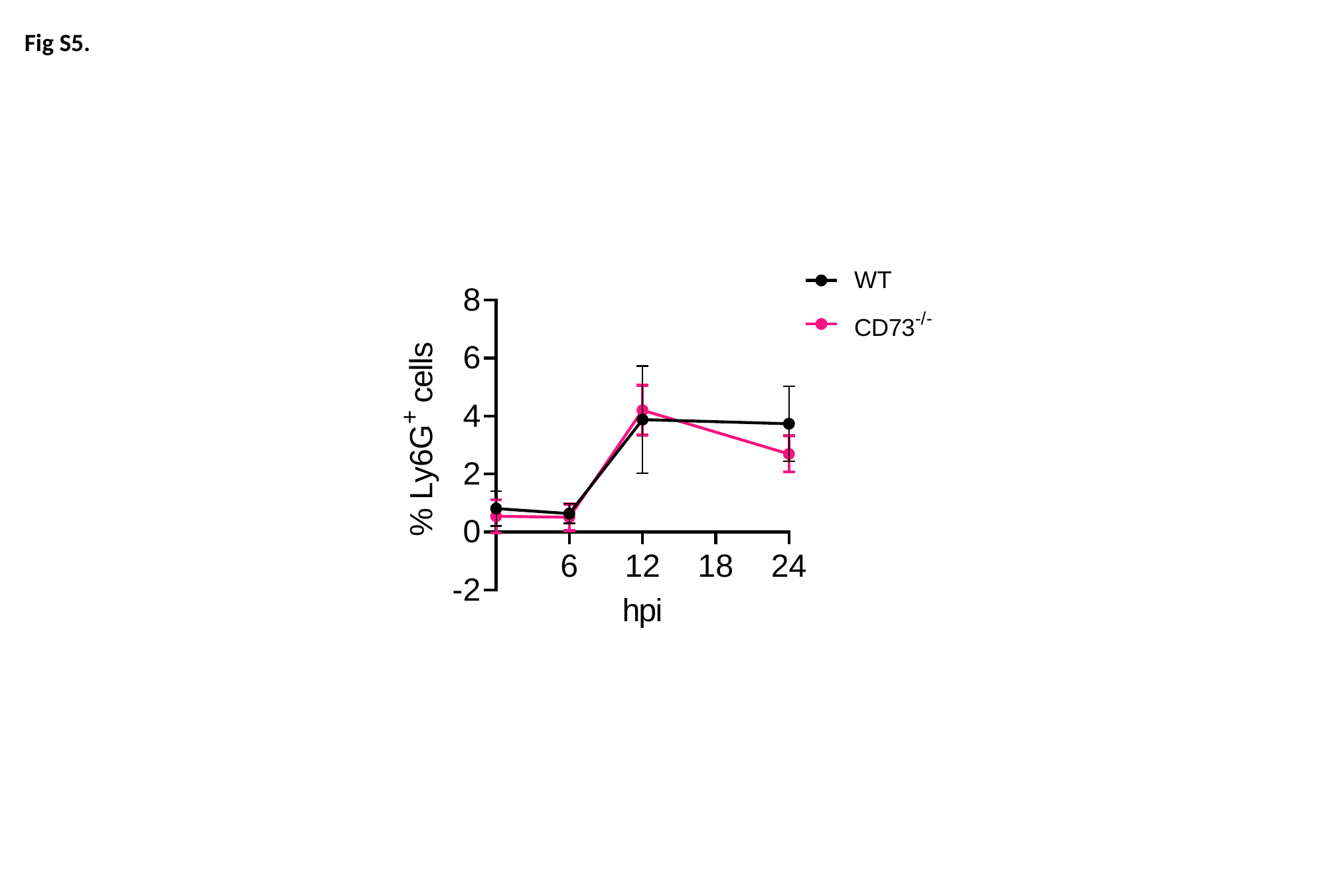

Fig S5.

### Slide 6
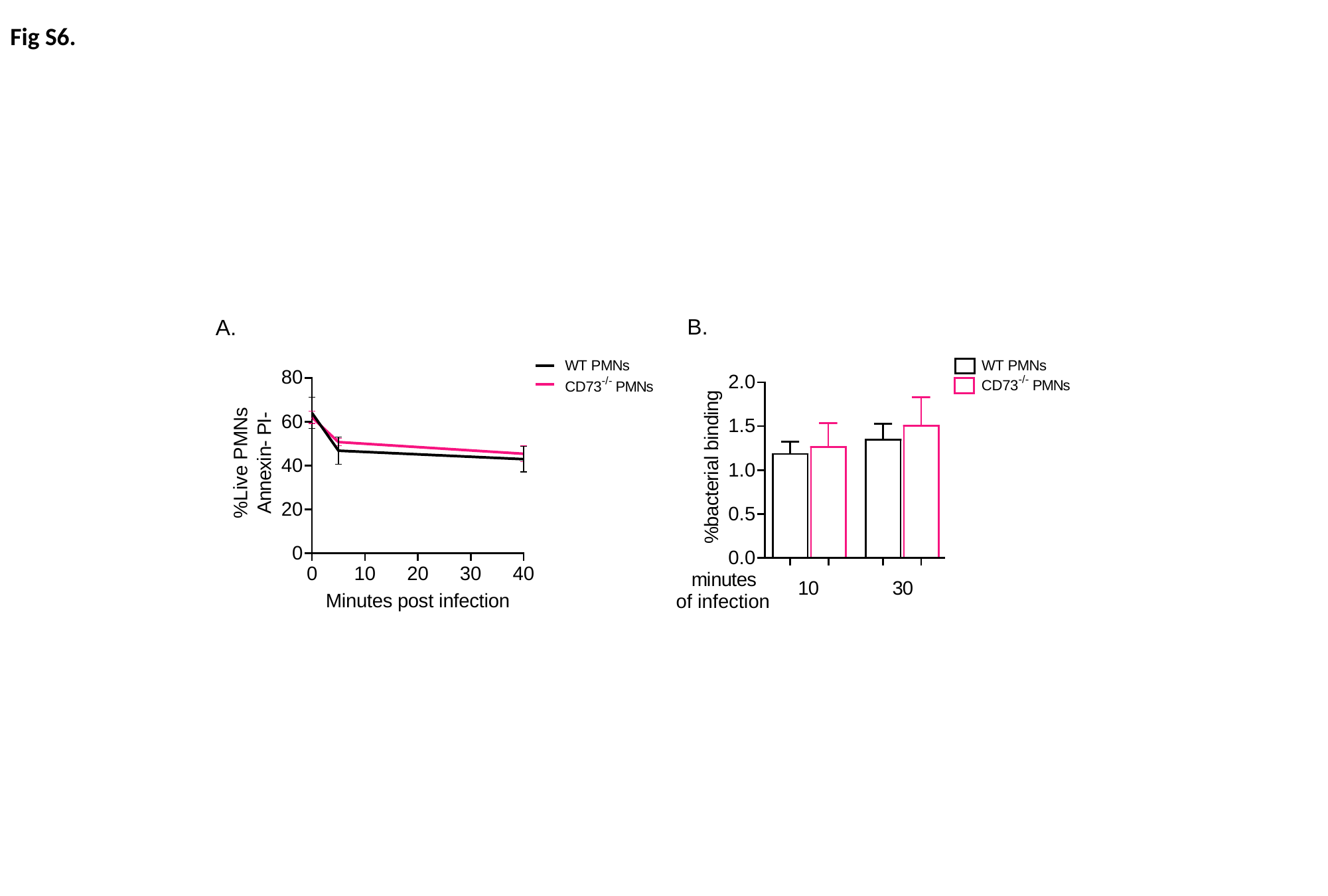

Fig S6.
B.
A.
