## Supplementary material for "Neutrophil-driven cardiac damage during invasive *Streptococcus pneumoniae* infection is regulated by CD73": Supplemental.docx

**Fig S1. Viable PMNs are observed in the cardiac tissue of infected mice.** WT (C57BL/6) mice were infected intra-peritoneally (I.P) with 10^3 CFU of *S*. *pneumoniae* TIGR4 and heart tissue was harvested at 6-, 12- and 24hpi to determine the viability of PMNs that influx into the cardiac tissue. Gating strategy showing PMN population (Ly6G^+^) in the heart at indicated time points under the live cell gate (non-fluorescent cells under the UV laser) and the dead cell gate (fluorescent cells under the UV laser) after cells were incubated with Live/Dead stain (Invitrogen) for 20 minutes.

**Fig S2. Bacterial burden in the cardiac tissue significantly correlates with the bacterial numbers in the blood.** Correlation analysis between bacterial burden in the blood and heart showing significant positive correlation between the two groups, as calculated by Pearson correlation analysis.

**Fig S3. Infection with *S. pneumoniae* D39 strain results in PMN influx in the host cardiac tissue.** WT (C57BL/6) mice were infected intra-peritoneally (I.P) with 10^3 CFU of *S*. *pneumoniae* D39 and heart tissue was harvested at 6-, 12- and 24hpi to detect PMN influx and bacterial presence. (A) Gating strategy showing PMN population (Ly6G^+^) in the heart at indicated time points (B) PMN enumeration in the heart (C) Bacterial load was enumerated in blood and (D) heart at 24hpi. Detection of PMNs in the cardiac tissue following D39 challenge was done with n = 3 mice in each time point. Bacterial enumeration in blood and heart was done with a total of 4 mice. Each circle represents one mouse. LOD: Limit of detection.

**Fig S4. Treatment of mice with anti-Ly6G antibodies does not affect the number of macrophages and dendritic cells in the cardiac tissue.** WT (C57BL/6) mice were injected intra-peritoneally (I.P) with either anti-Ly6G (clone 1A8) antibody or isotype control. At 18 hours later, the mice were injected with another dose of anti-Ly6G antibody and infected (I.P) with 10^3 CFU of *S. pneumoniae* TIGR4 or mock infected with PBS. The mice were euthanized 24hpi to harvest blood and heart. Flow cytometry was performed to confirm the depletion of PMNs (blood and heart) and enumerate macrophage and dendritic cell numbers. (A) Gating strategy showing PMN population (CD45^+^Gr-1^+^CD11b^+^), macrophages (CD45^+^F4/80^+^Ly6C^-^) and dendritic cells (CD45^+^F4/80^-^CD11c^+^). (B-D) Data showing the % PMNs in blood and heart (B), %macrophages in heart (C) and %dendritic cells in heart (D). Data shown are from a total of n = 4 mice per group.  Asterisks indicate significant differences as calculated by Student’s t-test with Welch’s correction with *p*<0.05 taken as significant.

**Fig S5.  CD73 does not play a role in PMN migration into the cardiac tissue.** C57BL/6 WT and CD73*^-/-^* mice were infected intra-peritoneally (I.P) with 10^3 CFU of *S. pneumoniae* TIGR4. The mice were then euthanized at 6-, 12- and 24hpi to harvest the hearts and quantify PMNs (Ly6G^+^) using flow cytometry. Data shown are pooled from a total of 6 mice per group.

**Fig S6. Bacterial binding to HL-1 cells is not affected by absence of CD73 on PMNs.**

Approximately 2 × 10^5^ PMNs from the bone-marrow of WT and CD73^-/-^ mice were infected with pre-opsonized *S. pneumoniae* TIGR4 (MOI of 5-10) at 37°C for the indicated time points or were mock treated with HBSS buffer containing 3% mouse serum (as un-infected condition) with and without the presence of HL-1 cardiomyocytes. (A) Flow cytometry was used to determine the percentages of live, apoptotic, or necrotic cells using the fluorescein isothiocyanate (FITC) annexin V apoptosis detection kit with propidium iodide (PI) (BioLegend) following the manufacturer’s protocol. (B) HL-1 cells following incubation with bacteria and PMNs were washed 5 times with PBS, trypsinized and plated on blood agar plates to enumerate bacterial numbers bound to the cells and %binding calculated with respect to the initial input. Graph in part (A) is a representative data from one of three independent experiments with two technical replicates per experiment (total n=6). Graph in part (B) is data collected from one experiment with three technical replicates.
